# Differential regulation of SHH signaling and the developmental control of species-specific jaw size through neural crest-mediated Gas1 expression

**DOI:** 10.1101/2021.12.17.473230

**Authors:** Zuzana Vavrušová, Daniel B. Chu, An Nguyen, Jennifer L. Fish, Richard A. Schneider

## Abstract

Developmental control of jaw size is crucial to prevent disease and facilitate evolution. We have shown that species-specific differences in jaw size are established by neural crest mesenchyme (NCM), which are the jaw progenitors that migrate into the mandibular primordia. NCM relies on multiple signaling molecules including Sonic Hedgehog (SHH) to mediate interactions with mandibular epithelium that facilitate outgrowth of the jaws. SHH signaling is known to promote outgrowth and so we tested if differential regulation of the SHH pathway can account for species-specific variation in mandibular primordia size. We analyze gene expression of SHH pathway members in duck, chick, and quail, and find higher transcriptional activation in the larger mandibular primordia of duck relative to those of chick and quail. We generate quail-duck chimeras and demonstrate that such activation is NCM-mediated. Gain- and loss-of-function experiments reveal a species-specific response to SHH signaling, with the target *Gas1* being most sensitive to manipulations. *Gas1* overexpression and knockdown in NCM alters cell number and/or mandibular primordia size. Our work suggests that NCM-mediated changes in SHH signaling may modulate jaw size during development, disease, and evolution.

**Summary Statement:** We have determined that *Gas1*, which is a component of the SHH signaling pathway, plays a key role in the development and evolution of jaw size.

## Introduction

Animals belonging to a given taxon typically share a conserved body plan that contains equivalent (i.e., homologous) structures. Such structures often appear to be iterative versions of a common template that can vary in relative size and/or proportions along a normal distribution, which is a phenomenon known as allometry or biological scaling (Thompson, 1917; Huxley, 1932; Huxley and Teissier, 1936; Woodger, 1945; Gould, 1966; Stern and Emlen, 1999; Gayon, 2000; Young et al., 2014; Schneider, 2018a). For purposes of functional morphology, the proper scaling of structures is robustly maintained during the development of individuals even though the size of these same structures can vary enormously both within a species and among members of related taxa (Russell, 1916; Haldane, 1926; Smith et al., 2015). Additionally, tissue regeneration and transplantation experiments indicate that structures retain intrinsic mechanisms enabling them to know their proper size and to regulate growth (Leevers and McNeill, 2005; Allard and Tabin, 2009; Fish et al., 2014; Uygur et al., 2016; Schneider, 2018b). But how these intrinsic mechanisms function and how they potentiate normal to abnormal phenotypic variation in size, is poorly understood. Moreover, molecular and cellular processes that enable early embryos to establish growth trajectories in support of requisite adult form and function remain to be identified especially as a means to understand etiologies of disease and mechanisms of evolution (Schneider, 2015; Woronowicz and Schneider, 2019).

On the molecular level, biological scaling likely involves species-specific modulations to intrinsic levels and patterns of gene expression that affect the behaviors of cells and the corresponding growth of tissues and organs. Such ideas emerged in the first half of the 20th century with the discovery of genes that alter the timing and rates of development (Huxley, 1932; Goldschmidt, 1938; Goldschmidt, 1940; de Beer, 1954), and they led to theories and quantitative methods during the rebirth of evolutionary developmental biology in the 1970s predicting how even small changes affecting developmental time and rates could generate significant phenotypic variation and transformations in size (Gould, 1977; Alberch et al., 1979; Hall, 1984; McKinney, 1988; Klingenberg, 1998; Smith, 2003; Keyte and Smith, 2014). In this context, molecules that act as morphogens seem to play a key role, especially when genetic changes alter their expression levels, source (i.e., cells that produce the morphogen), distribution (i.e., release of the morphogen), transportation (i.e., diffusion of the morphogen), and detection (i.e., cellular sensitivity to the morphogen) within the local environment (Oster et al., 1988; Gurdon et al., 1999; Gurdon and Bourillot, 2001; Dessaud et al., 2007; Tostevin et al., 2007; Ribes and Briscoe, 2009; Ben-Zvi and Barkai, 2010; Cheung et al., 2014). One well studied example is the classic morphogen Sonic Hedgehog (SHH), which elicits cellular proliferation or differentiation responses in a concentration-dependent manner that pattern and scale organs such as the limb bud, neural tube, and craniofacial primordia (Summerbell et al., 1973; Echelard et al., 1993; Riddle et al., 1993; Laufer et al., 1994; Ericson et al., 1995; López-Martínez et al., 1995; Ericson et al., 1997; Yang et al., 1997; Briscoe et al., 2001; Schneider et al., 2001; Zeng et al., 2001; Dessaud et al., 2007; Dessaud et al., 2008; McLellan et al., 2008; Hu and Marcucio, 2009; Young et al., 2010; Xu et al., 2015; Uygur et al., 2016).

SHH binds to the canonical receptor Patched (PTCH1) and the co-receptors Cell Adhesion Molecule-related/downregulated by Oncogenes (CDON) and Brother of CDON (BOC), as well as Growth Arrest-Specific 1 (GAS1) (Tenzen et al., 2006; Beachy et al., 2010; Izzi et al., 2011; Choudhry et al., 2014). The binding of PTCH1 by SHH results in de-repression and accumulation of Smoothened (SMO), and signal transduction through the Glioma-associated oncogene (GLI) transcription factors (van den Heuvel and Ingham, 1996; Alcedo and Noll, 1997; Quirk et al., 1997; Murone et al., 1999; Taipale et al., 2002; Wilson and Chuang, 2010). GLI2 and GLI3 are bifunctional transcription factors that can either activate or inhibit transcription whereas GLI1 lacks a transcriptional repressor domain and functions as a transcriptional activator (Marigo et al., 1996; Lee et al., 1997; Mo et al., 1997; Matise and Joyner, 1999; Wang et al., 2000; Bai and Joyner, 2001; Bai et al., 2002; Hui and Angers, 2011). The binding of CDON, BOC, and GAS1 by SHH promotes cell survival, proliferation, and differentiation, which ultimately can affect the size and shape of organs (Allen et al., 2007; Martinelli and Fan, 2007; Izzi et al., 2011; Delloye-Bourgeois et al., 2014).

In the current study, we tested if differential regulation of the SHH pathway can account for species-specific scaling of the developing jaw primordia by comparing expression of pathway members in duck, chick, and quail, which are three species of birds with distinctly different jaw sizes (i.e., from relatively large to small) and rates of maturation (Figure 1A, Supplemental Table 1). We analyze embryonic stages starting when neural crest mesenchyme (NCM), which are the jaw precursor cells (Le Lièvre, 1978; Noden, 1978; Jheon and Schneider, 2009), first arrive in the mandibular primordia and then undergo their patterned outgrowth. Previously we have shown that NCM controls species-specific jaw size (Schneider and Helms, 2003; Schneider, 2005; Schneider, 2015; Schneider, 2018b) and mechanisms contributing to the larger jaw size of duck versus quail include the allocation of approximately 15% more jaw progenitors to the mandibular primordia during early stages of migration and a longer cell cycle length (13.5 hours in duck and 11.0 hours in quail) (Fish et al., 2014). When taken alongside equivalent rates of proliferation over a period of 45 hours in duck versus 32 hours in quail (i.e., the amount of absolute time over comparable stages), this translates into a two-fold difference in cell number by embryonic stage (HH) 20. NCM also exerts species-specific control over multiple signaling pathways during later stages of cell proliferation and skeletal differentiation that directly affect jaw size (Eames and Schneider, 2008; Merrill et al., 2008; Fish and Schneider, 2014b; Hall et al., 2014; Ealba et al., 2015). Signaling interactions between NCM and the epithelium of the pharyngeal endoderm, which secretes SHH, promote the patterned outgrowth of the mandibular primordia and establish the anteroposterior polarity of the jaw skeleton (Couly et al., 2002; Helms and Schneider, 2003; Abzhanov and Tabin, 2004; Graham et al., 2005; Moore-Scott and Manley, 2005; Brito et al., 2006). Similarly, SHH is associated with species-specific shape and outgrowth of the face (Schneider et al., 2001; Young et al., 2010; Hu and Marcucio, 2012; Hu et al., 2015a; Hu et al., 2015b) and disruptions to SHH co-receptors as well as other pathway members can result in micrognathia and other jaw defects especially in association with holoprosencephaly (Mo et al., 1997; Cole and Krauss, 2003; Melnick et al., 2005; Zhang et al., 2006; Seppala et al., 2007; Ribeiro et al., 2010; Roessler and Muenke, 2010; Allen et al., 2011; Bae et al., 2011; Hui and Angers, 2011; Zhang et al., 2011; Dennis et al., 2012; Pineda-Alvarez et al., 2012; Seppala et al., 2014; Xavier et al., 2016; Hong et al., 2017; Echevarría-Andino and Allen, 2020). But the extent to which the SHH pathway is differentially regulated by NCM and whether changes to its regulation might affect the species-specific growth of the jaw primordia have not been tested.

**Figure 1.**
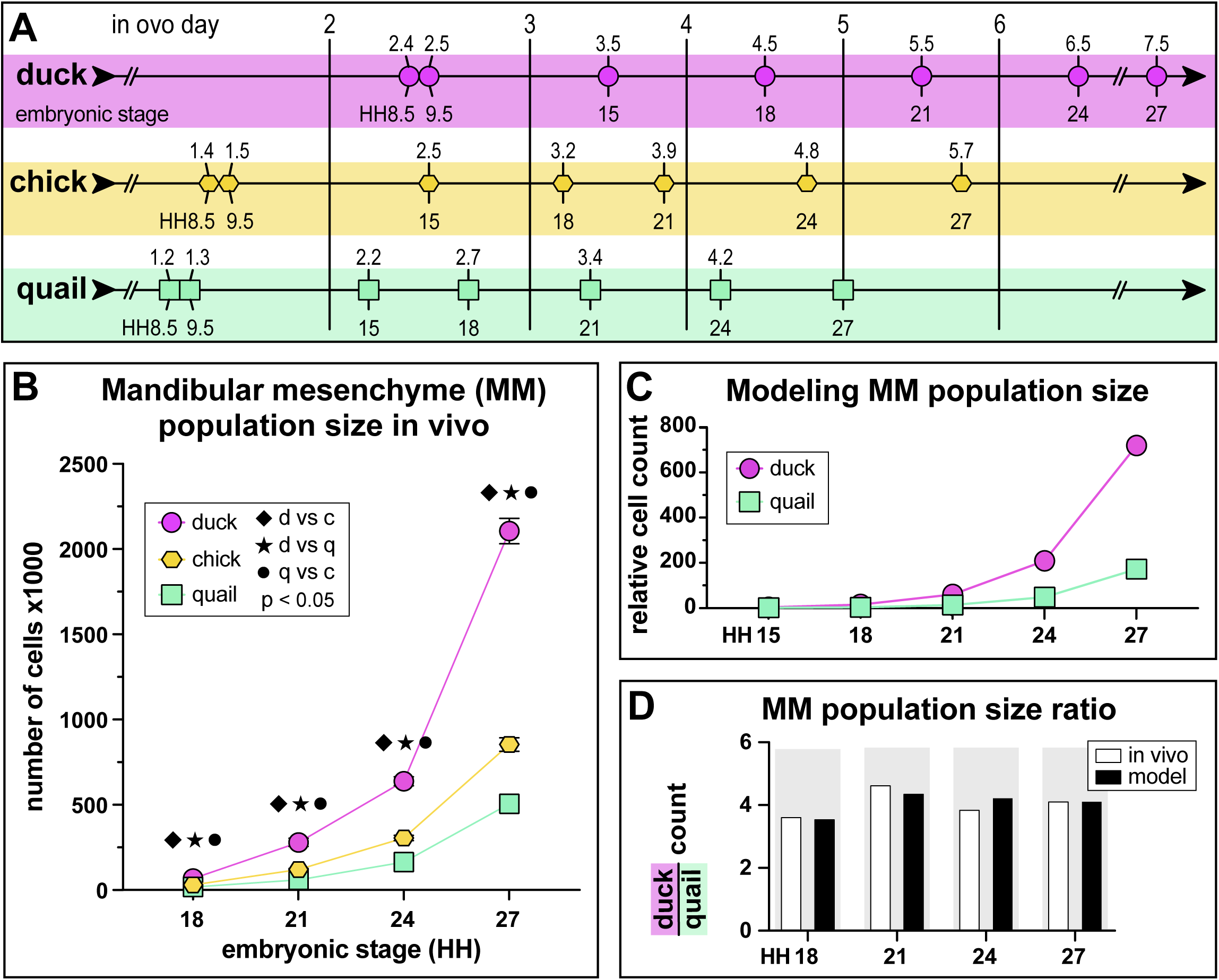
Species-specific variation at early embryonic stages. **(A)** The Hamburger-Hamilton (HH) staging system (i.e., embryonic stage) functions independent of time (i.e., absolute time), and instead relies on external morphological characters (i.e., developmental time). Thus, duck (violet 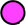), chick (yellow 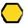), and quail (green 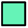) embryos can be aligned at equivalent stages by incubating them for different lengths of time. *In ovo* day represents the number of incubation days needed to reach equivalent stages for each species. **(B)** Mandibular mesenchyme (MM) population size at HH18, HH21, HH24, and HH27 in duck (violet), chick (yellow), and quail (green) embryos. **(C)** Relative mandibular mesenchyme population size modelling at HH15, HH18, HH21, HH24, and HH27 in duck (violet) and quail (green) embryos using the number of migratory NCM cells, cell cycle length, and absolute time. **(D)** Relative mandibular mesenchyme population size based on duck to quail ratios for in vivo (white) and model (black) data at HH18, HH21, HH24, and HH27.

We performed a comparative analysis of the developing mandibular primordia of duck, chick, and quail from HH15 to HH27 and find species-specific differences in the amount of mandibular mesenchyme at each stage. We quantified expression of *Shh*, *Ptch1*, *Cdon*, *Boc*, *Gas1*, *Smo*, *Gli1*, *Gli2*, and *Gli3* in mandibular primordia of duck, chick, quail, and chimeras in which we transplanted presumptive NCM from quail to duck (i.e., quck). Quck chimeras offer an effective way to assess the extent to which NCM differentially regulates gene expression due to the faster maturation rate of the quail donor versus the duck host (Schneider and Helms, 2003; Eames and Schneider, 2005; Merrill et al., 2008; Tokita and Schneider, 2009; Solem et al., 2011; Hall et al., 2014; Ealba et al., 2015; Woronowicz et al., 2018). Our strategy uncovers stage- and species-specific expression levels for some key pathway members but not others, identifies stage- and species-specific levels of pathway activation, and reveals that these differences are NCM-mediated since donor NCM maintains its quail-like expression in duck hosts. We also tested if duck, chick, and quail have an intrinsic species-specific response to SHH signaling by culturing explants of mandibular primordia and treating them with different concentrations of recombinant SHH (rSHH) protein or SHH pathway inhibitor. These pathway activation and inhibition experiments reveal a species-specific response to SHH signaling, with the SHH co-receptor *Gas1* being most sensitive to manipulations. This is in contrast to *in vitro* studies that we perform in parallel where we observe a minimal response to the same treatments. Finally, *in ovo* overexpression and knockdown of *Gas1* in NCM alters cell number and/or mandible size. Overall, our work suggests that species-specific changes in the response of NCM to SHH signaling and the differential regulation of *Gas1* expression may be a mechanism through which NCM controls jaw size during development, disease, and evolution.

## Results

### Species-specific differences arise early and progress during development

To compare the size of mandibular primordia in duck, chick, and quail embryos, we quantified the number of mandibular mesenchymal cells in HH18, HH21, HH24, and HH27 embryos (Figure 1B). We observe significant differences in mandibular mesenchyme population size among all three species for every embryonic stage (n and p-values in Supplemental Tables 2 and 3). Mandibular mesenchyme population size is significantly larger in duck and chick than in quail throughout the developmental stages with duck being the largest.

To evaluate the rate at which relative mandibular mesenchyme population size increases over time in duck and quail we calculated duck to quail mandibular mesenchyme population size ratios at HH18, HH21, HH24, and HH27. The mandibular mesenchyme population size ratio is lowest at HH18 and highest at HH21 with very small changes over development (values in Supplemental Table 4). The mandibular mesenchyme population is on average 4.1-fold larger in duck than quail.

To determine whether differences in the mandibular mesenchyme population size between duck and quail are due to differences in cell cycle length, initial size of the migratory NCM population (Fish et al., 2014), and/or the absolute time that each species takes to reach each successive developmental stage (Figure 1A) we modeled relative mandibular mesenchyme population size, using the number of migratory NCM, cell cycle length, and absolute time (Figure 1C, Supplemental Table 5). We compared modeled data to the population size *in vivo* (Figure 1D, Supplemental Table 4) and we were able to replicate the relative mandibular mesenchyme population size observed *in vivo* for duck and quail. When accounting for differences in cell cycle length (i.e., proliferation) and developmental rate (i.e., absolute time), we observe an approximately 26% increase in the size of the pre-migratory NCM population in duck compared to quail embryos (Fish et al., 2014), which leads to an average 4.1-fold larger population of mandibular mesenchyme in duck.

### SHH pathway activation is species-specific

Prior studies have shown that SHH regulates growth and proliferation of the mandibular primordia (Brito et al., 2008; Roper et al., 2009; ten Berge et al., 2001) and that *Shh* is expressed in similar domains (i.e., in the pharyngeal endoderm) in duck, chick, and quail embryos (Schneider et al., 2001; Fish et al., 2014). To test the hypothesis that species-specific expression of SHH pathway members underlies the differential rates of proliferation that ultimately affect mandibular mesenchyme population size, we compared the expression of SHH pathway members and targets among duck, chick, and quail mandibular primordia during development. Using qPCR we quantified the relative expression of key components including *Shh*, *Ptch1*, *Gas1*, *Gli1* (Figure 2A to 2D), *Boc*, *Cdon*, *Smo*, *Gli2*, and *Gli3* (Supplemental Figure 1) in duck, chick, and quail mandibular primordia at HH15, HH18, HH21, HH24, and HH27. We find stage- and species-specific differences in many genes with the most striking difference in the relative expression of *Gas1*. *Gas1* levels are approximately 25 to 75 times higher in duck than in chick and quail with these differences increasing from HH15 to HH27 (Figure 2C; for all significant comparisons between species p < 0.02, and n and p-values are listed in Supplemental Tables 6 and 8).

**Figure 2.**
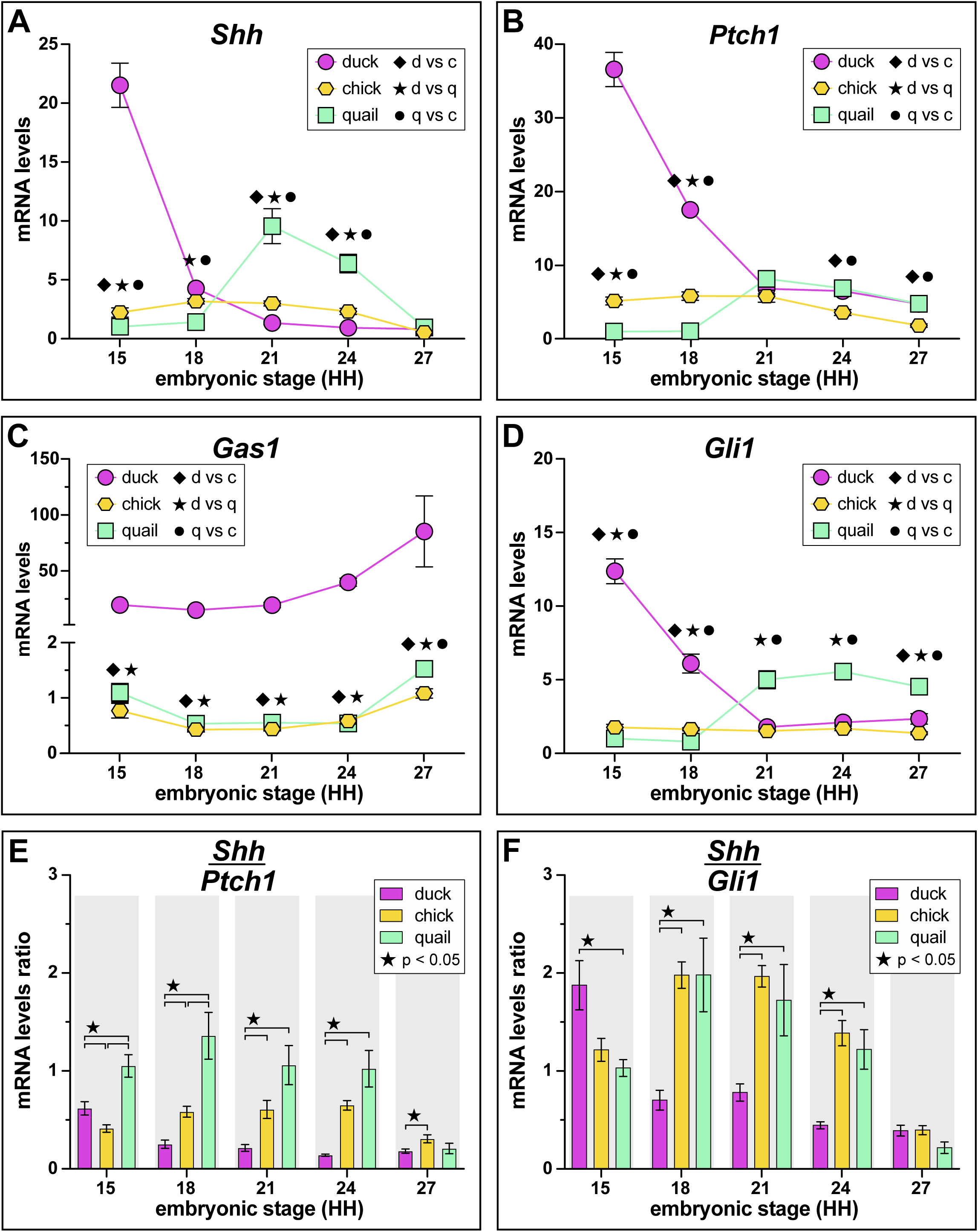
Expression of SHH pathway members and pathway activation in mandibular primordia of duck, chick, quail, and chimeric quck. Relative mRNA levels of **(A)** *Shh*, **(B)** *Ptch1*, **(C)** *Gas1*, and **(D)** *Gli1* in mandibular primordia of duck (violet), chick (yellow), and quail (green) embryos at HH15, HH18, HH21, HH24, and HH27. Expression levels were assayed by qRT-PCR and normalized to *r18s*. Significance is shown (p-value < 0.02, n ≥ 6 for each group and data point, and error bars represent SEM) for comparisons between different species at the same embryonic stage (i.e., diamond symbol for duck versus chick, asterisk symbol for duck versus quail, and full circle symbol for quail versus chick). SHH pathway activation is represented by ratios of **(E)** *Shh* to *Ptch1* and **(F)** *Shh* to *Gli1* mRNA levels for duck, chick, and quail at HH15, HH18, HH21, HH24, and HH27. Significance is shown for comparisons between species as denoted by brackets and p-values are as indicated. n ≥ 6 for each group and data point, and error bars represent SEM.

To assess if there are species-specific differences in the transcriptional activation of the SHH pathway among duck, chick, and quail mandibular primordia, we calculated gene expression ratios for *Shh* to *Ptch1* (Figure 2E) and *Shh* to *Gli1* (Figure 2F) at HH15, HH18, HH21, HH24, and HH27. For *Shh* to *Ptch1* ratios we observe significantly higher transcriptional activation in duck from HH15 to HH24 compared to quail, and from HH18 to HH27 compared to chick, with quail having the lowest pathway activation. For *Shh* to *Gli1* ratios we observe significantly higher transcriptional activation for chick and quail at HH15 compared to duck and this becomes significantly higher in duck from HH18 to HH24 (n and p-values in Supplemental Tables 7 and 8).

### NCM differentially regulates the SHH pathway at multiple levels

To assess the extent to which the SHH pathway is differentially regulated by NCM, we compared gene expression in the mandibular primordia of chimeric quck to that observed in duck and quail (Figure 3A). We quantified relative expression of *Shh*, *Boc*, *Cdon*, *Gas1*, *Smo*, *Gli1* (Figure 3B to 3G), *Ptch1*, *Gli2*, and *Gli3* (Supplemental Table 6) in duck, quail, and quck mandibular primordia at quck embryonic stage HH21 and HH24 (i.e., host cells at HH21 and HH24; donor cells at HH24 and HH27 respectively). We find NCM-mediated gene expression for *Gas1* and *Gli1*. We observe significantly lower gene expression in quck relative to host (i.e., duck) for *Gas1* (Figure 3E) and significantly higher expression for *Gli1* (Figure 3G) in quck compared to host at both embryonic stages, while expression of *Shh*, *Boc*, *Cdon*, and *Smo* remains host-like (i.e., duck-like).

**Figure 3.**
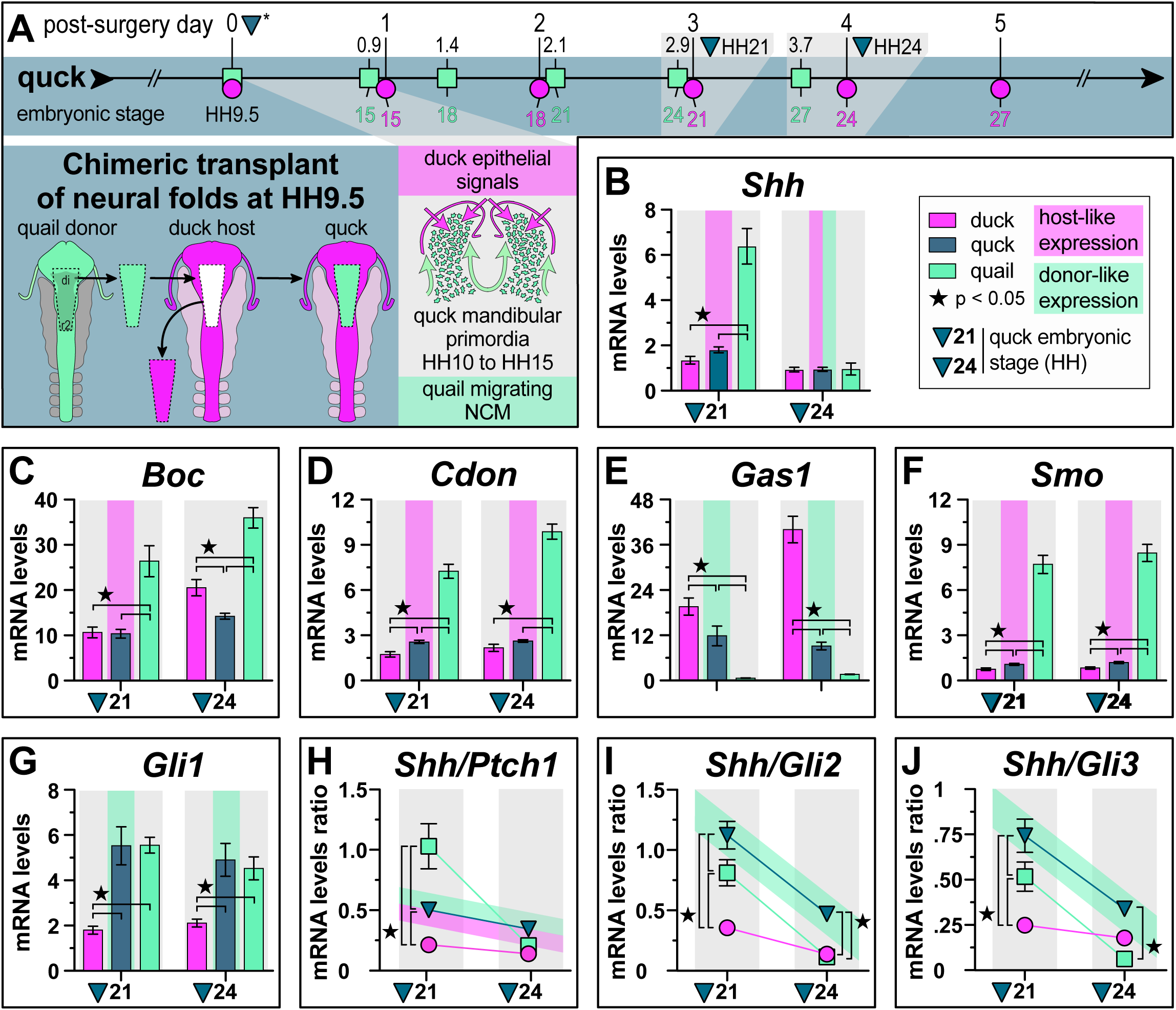
Neural-crest mediated gene expression of SHH pathway members in mandibular primordia. **(A)** To generate quail-duck chimeras (“quck”), quail and duck embryos are stage-matched for surgery at HH9.5 (i.e., post-surgery day 0, 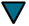*) by starting the incubation of their eggs at different times (see Figure1A). Bilateral neural folds from the mid-diencephalon (di) to rhombomere 2 (r2) of the hindbrain (dark green), which generate neural crest mesenchyme (NCM), are transplanted from quail to duck. Quail donor NCM (green cells) migrates (green arrows) into mandibular primordia between HH10 and HH15. Due to its faster rate of maturation, quail NCM develops approximately three stages ahead of the slower-maturing duck host within two days post-surgery. Quail NCM receives cues from and interacts with duck-host derived epithelium (violet arrows). Relative mRNA levels of **(B)** *Shh*, **(C)** *Boc*, **(D)** *Cdon*, **(E)** *Gas1*, **(F)** *Smo*, and **(G)** *Gli1* in mandibular primordia of duck (violet), quck (blue), and quail (green) embryos at quck embryonic stage HH21 (i.e., host-duck cells at HH21 and donor-quail cells at HH24) and HH24 (i.e., host-duck cells at HH24 and donor-quail cells at HH27). Gene expression ratios of relative mRNA levels of **(H)** *Shh* to *Ptch1*, **(I)** *Shh* to *Gli2*, and **(J)** *Shh* to *Gli3* in mandibular primordia of duck, quck, and quail at quck embryonic stage HH21 and HH24. Expression levels were assayed by qRT-PCR and normalized to *r18s*. Significance is shown for comparisons between species as denoted by brackets and p-values are as indicated. n ≥ 6 for each group and data point, and error bars represent SEM. Host-like expression is denoted where the relative mRNA expression and/or the expression pattern is not significantly different and/or similar to host (i.e., duck) values. Donor-like expression represents when the relative mRNA expression and/or the expression pattern is significantly different fron the host (i.e., duck) and/or similar to donor (i.e., quail) values.

To assess the effects of NCM on SHH pathway activation, we calculated gene expression ratios for *Shh* to *Ptch1.* We also assessed the effects of NCM on the ratios of *Shh* to *Gli2* and to *Gli3* (Figure 3H to 3J). For *Shh* to *Ptch1,* we observe significantly lower quail-like pathway activation (i.e., lower *Ptch1* expression) in quck at both embryonic stages while the pattern of change in the ratio over time remains duck-like (Figure 3H). For *Shh* to *Gli2* and *Shh* to *Gli3* we observe significantly higher ratios for quck compared to duck at both embryonic stages, and find that the patterns of change in the ratio over time are more quail-like (i.e., donor-like) (Figure 3I to 3J; n and p-values are in Supplemental Tables 7 and 11). Thus, we find that NCM regulates absolute levels of expression for some pathway members (i.e., *Gli1*, Figure 3G), causes intermediate changes to gene expression levels (i.e., *Gas1* and *Ptch1*, Figure 3E, 3H), and alters the pattern of change in the ratio over time (i.e., *Shh* to *Gli2* and to *Gli3*, Figure 3I to 3J).

### SHH pathway activation is dose-dependent

To determine if there are intrinsic species-specific differences in sensitivity to SHH signaling (i.e., a dose-response to pathway activator and/or inhibitor), we cultured explants of HH21 mandibular primordia from duck, chick, and quail and treated them with rSHH protein (i.e., pathway activator) or cyclopamine (i.e., pathway inhibitor). After 24 hours of treatment, we quantified relative expression of *Ptch1*, *Gas1*, *Gli1*, *Gli2*, *Gli3* (Figure 4), *Shh*, *Boc*, *Cdon*, and *Smo* (Supplemental Figure 2). We find that duck and chick show a greater response to rSHH treatments compared to quail for both *Ptch1* (Figure 4A) and *Gli1* (Figure 4C) whereas the response of these genes to cyclopamine is similar in all three species. Most SHH pathway members exhibit species-specific differences in their relative expression except for *Shh* (Supplemental Figure 2A) and *Smo* (Supplemental Figure 2D) for which we observe no significant differences in relative expression between control and any treatment group for any species. Notably, we observe the greatest interspecies sensitivity to SHH pathway manipulations with *Gas1* (Figure 4B). *Gas1* expression is significantly downregulated starting at 0.1 ng/mL of SHH protein for quail, and at 100 ng/mL for chick compared to controls, whereas there is no significant change at any concentration of rSHH for duck. By contrast, *Gas1* expression is significantly increased in duck and chick when treated with cyclopamine, while there is no significant difference in quail (n and p-values for all genes and species are in Supplemental Tables 9 and 10).

**Figure 4.**
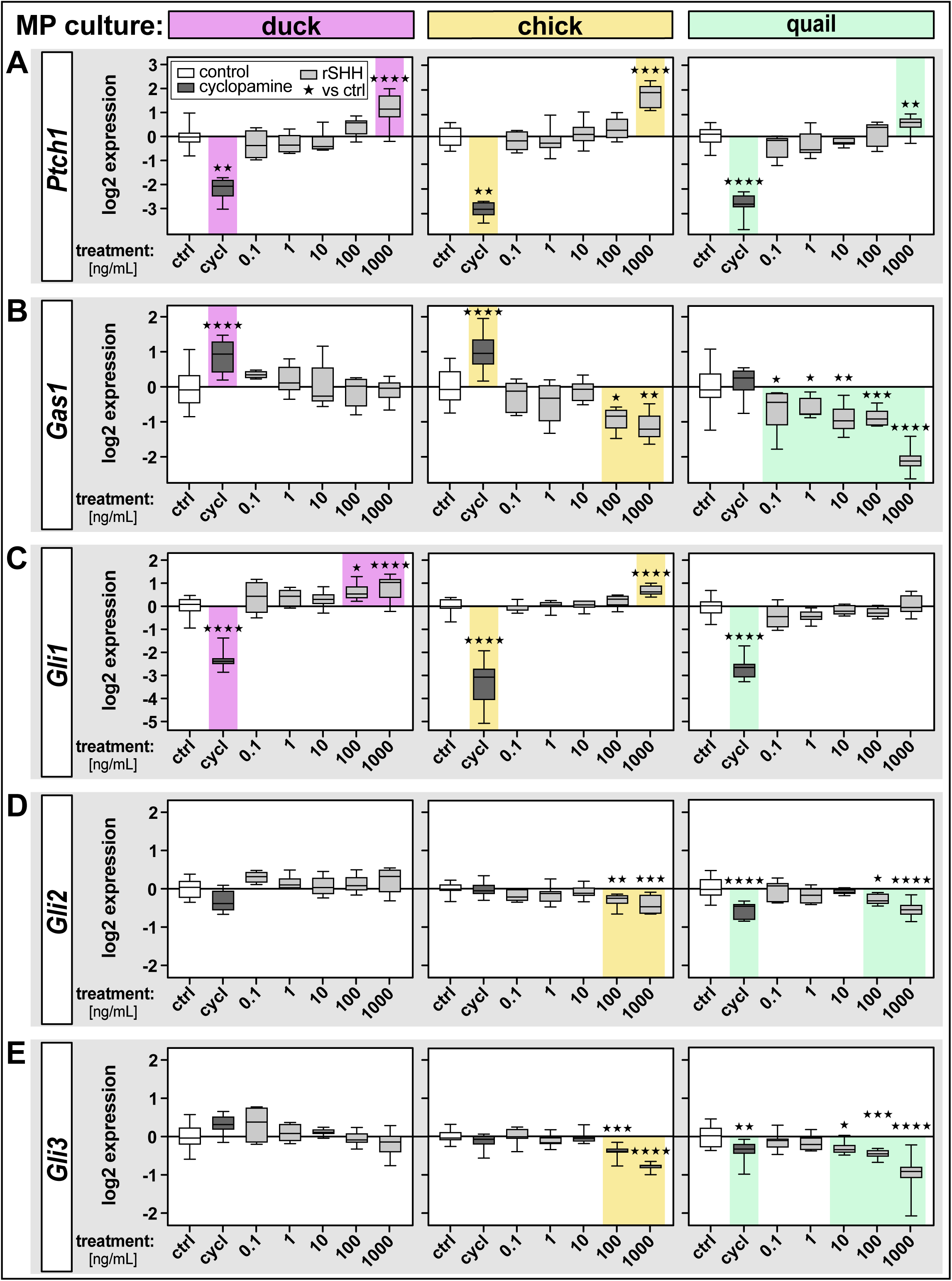
Effects of inhibition and activation of the SHH pathway in mandibular primordia of duck, chick, and quail. Mandibular primordia (MP) from duck (violet), chick (yellow), and quail (green) embryos were harvested at HH21, placed in culture, and treated with cyclopamine (cycl, dark grey) to inhibit the SHH pathway or with five concentrations of recombinant (r) SHH protein (light grey) to activate the SHH pathway. Box plots show relative levels of mRNA expression compared to controls (on the y-axis in log_2_ scale) for SHH pathway members including **(A)** *Ptch1*, **(B)** *Gas1*, **(C)** *Gli1*, **(D)** *Gli2*, and **(E)** *Gli3* 24 hours after treatment. Control (ctrl, white) and treatment groups are shown on the x-axis. Expression levels were assayed by qRT-PCR and normalized to *r18s*. Significance is shown for comparisons between control and treatment groups (i.e., cyclopamine or rSHH) within the same species as indicated by colored shading and by the following symbols for p-values: * p < 0.05, ** p < 0.01, *** p < 0.001, and **** p < 0.0001. p-values ≥ 0.05 are considered not significant (ns). n ≥ 4 for each group and data point.

### Spatial domains of Gas1 expression are conserved among duck, chick, and quail

To understand the spatial expression of *Gas1*, we performed whole mount *in situ* hybridization on HH21 and HH27 duck, chick, and quail mandibular primordia (Figure 5A to 5F). We find that *Gas1* expression is symmetrical for all three species with one lateral domain on each side of the mandibular primordia at HH21 and two distinct domains at HH27 (one lateral domain on each side and one medial domain). We also quantified *Gas1* expression in the mandibular mesenchyme versus epithelium of duck and chick at HH24 and HH27 (Figure 5G to 5H). We observed significantly higher *Gas1* expression in mandibular mesenchyme compared to epithelia for both species at both embryonic stages.

**Figure 5.**
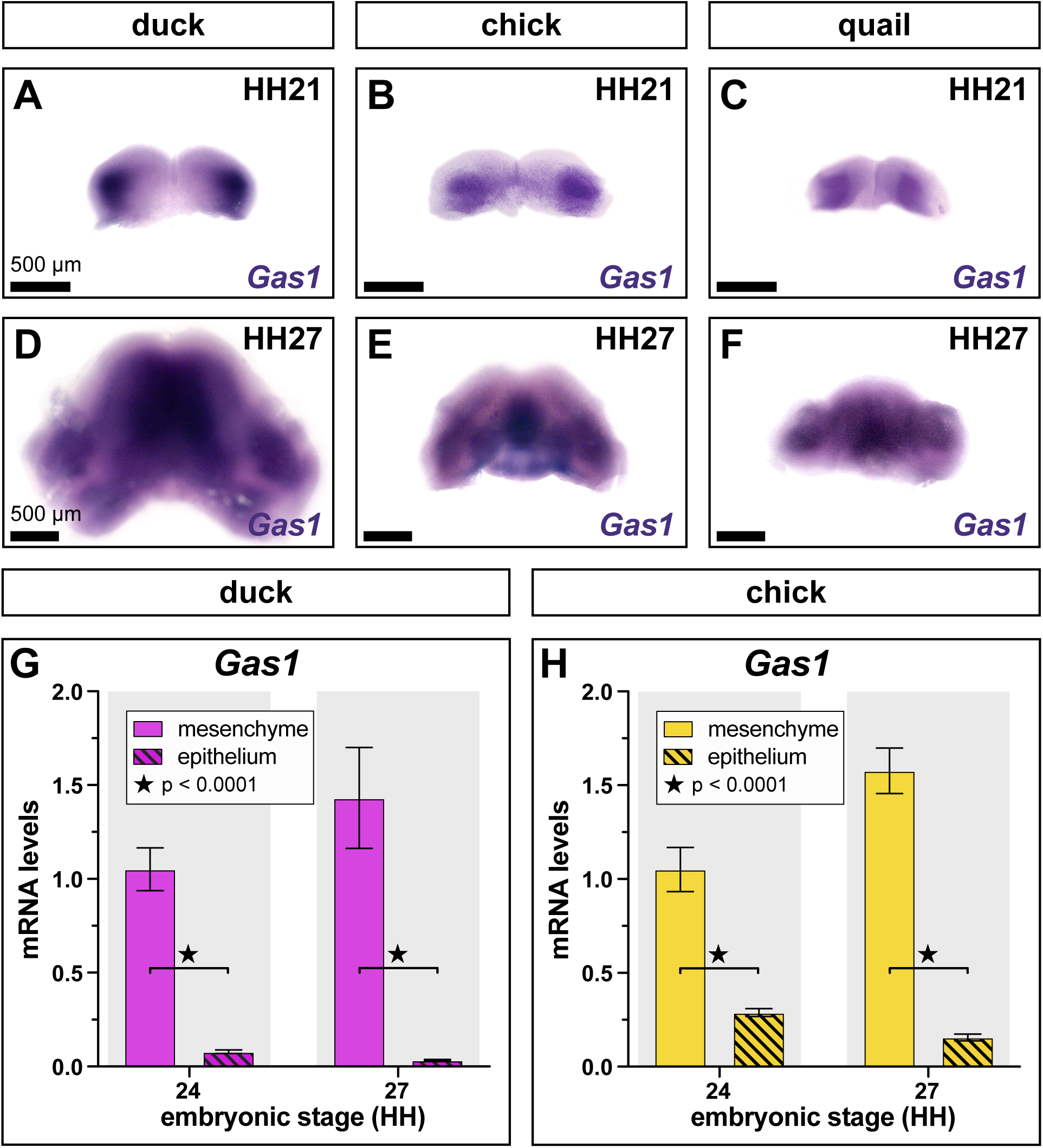
Expression of *Gas1* in mandibular primordia at early developmental stages. Whole mount *in situ* hybridization showing *Gas1* expression (purple stain) in mandibular primordia from **(A)** duck, **(B)** chick, and **(C)** quail at HH21; and in **(D)** duck, **(E)** chick, and **(F)** quail at HH27. **(G)** Relative levels of mRNA expression for *Gas1* in duck mandibular mesenchyme (violet) and epithelium (striped) at HH24 and HH27. **(H)** Relative levels of mRNA expression for *Gas1* in chick mandibular mesenchyme (yellow) and epithelium (striped) at HH24 and HH27. Expression levels were assayed by qRT-PCR and normalized to *r18s*. Significance is shown for comparisons between mesenchyme and epithelium at the same embryonic stage within species as denoted by brackets and p-values are as indicated. n ≥ 7 for each group and data point, and error bars represent SEM.

### Response to SHH treatments is context dependent

To test if the response to changes in SHH pathway activation is mediated by the context of the mandibular primordia or is instead a cell-autonomous effect, we treated chick fibroblasts (i.e., DF-1 cells) with the same varying concentrations of rSHH protein as mandibular primordia explant cultures. While treating with rSHH for 24 hours results in SHH pathway activation in chick fibroblasts (i.e., *Ptch1* expression significantly increased at 1000 ng/mL), there is no significant difference in *Gli1* or *Gas1* expression compared to control (i.e., 0 ng/mL of rSHH) (Figure 6A). Thus, treating chick fibroblasts with rSHH protein is not sufficient to alter expression and suggests that the response to SHH signaling is conferred by the developmental context of the mandibular primordia (n and p-values for all genes and treatments are in Supplemental Tables 12 and 13).

**Figure 6.**
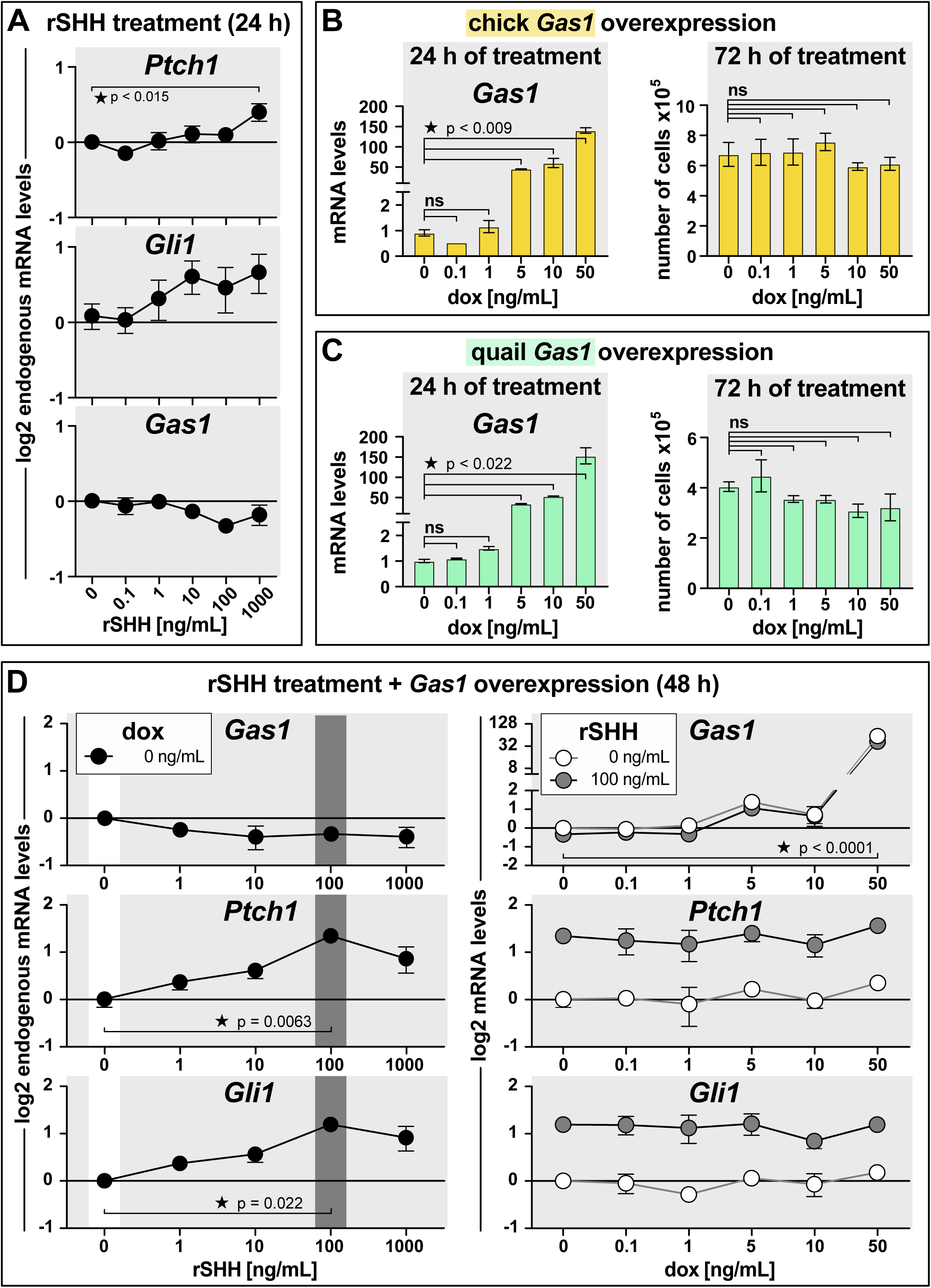
Effects of SHH pathway activation and *Gas1* overexpression in cell culture. **(A)** Relative levels of mRNA expression (on the y-axis in log_2_ scale) for *Gas1*, *Ptch1*, and *Gli1* in chick fibroblasts (i.e., DF-1) 24 hours (h) after treatment with five concentrations of recombinant (r) SHH protein. **(B)** Relative *Gas1* mRNA levels in chick fibroblasts 24 hours after doxycycline (dox)-induction of a stably integrated chick *Gas1* overexpression vector. Number of chick *Gas1*-positive cells 72 hours after induction with dox. **(C)** Relative *Gas1* mRNA levels in chick fibroblasts 24 hours after dox-induction of a stably integrated quail *Gas1* overexpression vector. Number of quail *Gas1*-positive cells 72 hours after induction with dox. **(D)** Relative levels of mRNA expression for *Gas1*, *Ptch1*, and *Gli1* in chick fibroblasts 48 hours after treatment with rSHH protein. Relative levels of mRNA expression for *Gas1*, *Ptch1*, and *Gli1* in chick fibroblasts 48 hours after dox-induction of a stably integrated *Gas1* overexpression vector and treatment with rSHH protein (control 0 ng/mL in white and 100 ng/mL dark grey). Expression levels were assayed by qRT-PCR and normalized to *r18s*. Significance is shown for comparisons between control and treatment groups as denoted by brackets and p-values are as indicated. P-values ≥ 0.05 are considered not significant (ns). n ≥ 2 for each group and data point, and error bars represent SEM.

To determine if variation in the levels of *Gas1* expression can affect expression of SHH pathway members and cell number, we overexpressed chick and quail *Gas1* in chick fibroblasts by using a doxycycline (dox)-inducible promoter system that have we characterized previously (Chu et al., 2020) and by varying the concentrations of dox. We confirmed chick (Figure 6B, left panel) and quail (Figure 6C, left panel) *Gas1* overexpression by qRT-PCR after 24 hours of dox treatment, and we quantified expression of *Shh* (data not shown, gene not expressed), *Boc*, *Cdon*, *Smo*, *Ptch1*, *Gli1*, *Gli2*, and *Gli3* (Supplemental Figure 3A). We do not observe any changes to the expression of these genes with the exception of *Smo,* which becomes elevated when quail *Gas1* is overexpressed using 0.1 ng/mL of dox. We also quantified the number of cells per plate after 72 hours of dox-induction (Figure 6B to 6C, right panel). Our results show no difference in the number of cells per plate in any of the treatment groups for either chick or quail *Gas1* overexpression compared to control (i.e., 0 ng/mL of dox). Thus, *Gas1* overexpression in chick fibroblasts does not affect gene expression of SHH pathway members 24 hours after dox-induction nor alter the number of cells per plate after 72 hours (n and p-values for all genes and treatments are in Supplemental Tables 14 to 16).

To evaluate if there is a relationship between *Gas1* overexpression and SHH pathway activation, we treated *Gas1*-expressing chick fibroblasts with five different concentrations of dox and four different concentrations of rSHH (Supplemental Table 17). With no dox treatment, we did not observe any change to *Gas1* expression at any rSHH concentration (Figure 6D, left panel). *Ptch1* and *Gli1* expression were significantly higher at 100 ng/mL rSHH, which confirms that the SHH pathway was activated. With *Gas1* overexpression, there is no significant difference in gene expression levels for *Ptch1*, *Gli1* (Figure 6D, right panel), *Boc*, *Cdon*, *Smo*, *Gli2*, and *Gli3* (Supplemental Figure 3B) following any dox treatment with the exception of *Cdon* at rSHH at 0 ng/mL and 10 ng/mL of dox. In addition, for samples at 1, 10, and 1000 ng/mL of rSHH we observed no significant difference for any of these genes at any dox concentration with the exception of *Gas1,* which is significantly higher for all three rSHH concentrations at 50 ng/mL of dox (data not shown). Thus, *Gas1* overexpression, while able to activate the SHH pathway, is not sufficient to alter the response of members of the SHH pathway in chick fibroblasts (n and p-values for all genes and treatments are in Supplemental Tables 17 and 18). Taken together, these data suggest that the species-specific response to SHH in mandibular primordia is context dependent.

### Changes in Gas1 expression affect mandibular primordia size in vivo

Our *in vivo* molecular data together with results from our SHH pathway manipulations in mandibular explant cultures suggest that *Gas1* is key component of the SHH pathway that may mediate species-specific differences in cell number and jaw size. To test if changes in levels of *Gas1* expression can affect the size of the mandibular primordia, we altered *Gas1* expression in duck and quail. First, we electroporated our dox-inducible *Gas1* overexpression plasmid bilaterally into the presumptive NCM of duck and quail embryos at HH8.5 and then treated these embryos with dox at HH15. We collected duck and quail embryos at HH18, HH21, and HH24 and quantified the number of mandibular mesenchymal cells. We observe a significant reduction in the amount of mandibular mesenchyme at HH21 and HH24 in duck embryos overexpressing *Gas1* (Figure 7A). The extent of *Gas1*-positive NCM was validated in whole mount by mScarlet fluorescence (RFP) at HH18 (Figure 7B and 7C), HH21 (Figure 7D and 7E), and HH24 (Figure 7F and 7G). Although not significant (p < 0.0614), we observed a trend towards reduction in the amount of mandibular mesenchyme in quail embryos (Figure 7H; n and p-values for all samples are in Supplemental Tables 3 and 19). To assess the effects of overexpression on mandibular morphology, we also electroporated *Gas1* overexpression plasmid unilaterally into the right side of presumptive NCM of quail embryos at HH8.5 and then treated these embryos with dox at HH15. By HH27, quail embryos show a clear reduction in the size of the developing jaw on the electroporated side (Figure 7I to 7L). Again, the extent of *Gas1*-positive overexpression in NCM in whole mount was validated by RFP in the treated versus internal control sides (Figure 7J to 7L).

**Figure 7.**
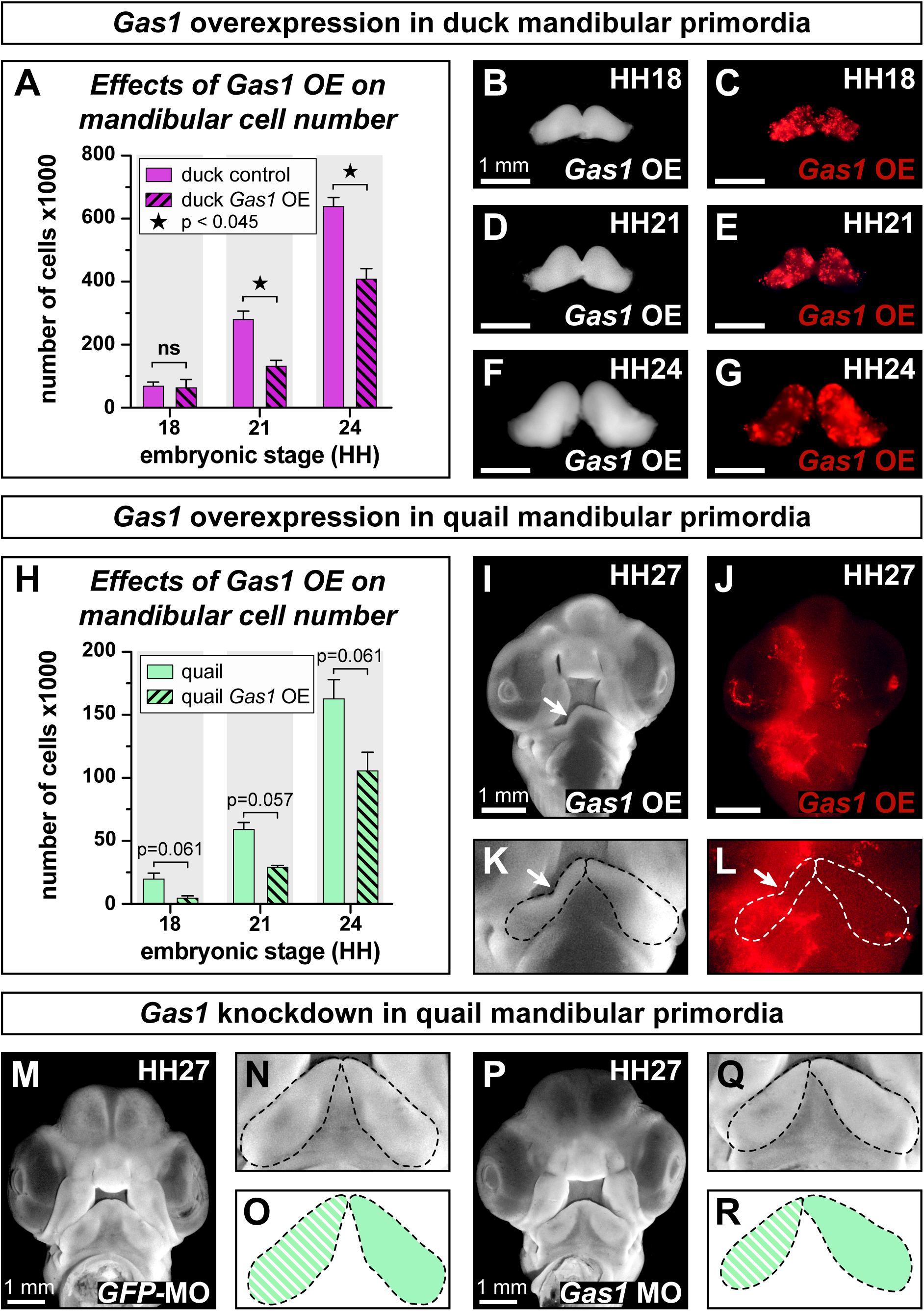
Effects of *Gas1* overexpression and knockdown in duck and quail embryos. **(A)** Mesenchymal cell number in duck mandibular primordia following bilateral *in ovo* electroporation of a stably integrating and doxycycline (dox)-inducible *Gas1* overexpression (OE) vector into NCM at HH8.5. *Gas1* overexpression was induced with 50 ng/mL of dox at HH15 and mandibular primordia were collected at HH18, HH21, and HH24. **(B)** Dissected duck mandibular primordia in whole mount at HH18 following bilateral *in ovo* electroporation of a *Gas1* overexpression vector into NCM at HH8.5. **(C)** *Gas1*-positive NCM can be visualized by mScarlet fluorescence (RFP). *Gas1* overexpression **(D)** in whole mount can be visualized **(E)** by RFP at HH21, and **(F)** in whole mount and **(G)** by RFP at HH24. **(H)** Mesenchymal cell number in quail mandibular primordia following bilateral *in ovo* electroporation of a *Gas1* overexpression vector into NCM at HH8.5. *Gas1* overexpression was induced with dox at HH15 and mandibular primordia were collected at HH18, HH21, and HH24. **(I)** Quail embryo in whole mount at HH27 following unilateral *in ovo* electroporation of a *Gas1* overexpression vector into NCM at HH8.5 and induction with dox at HH15. The treated size appears smaller (white arrow) than the contralateral control side. **(J)** Unilateral distribution of *Gas1*-positive NCM can be visualized by RFP on one side of the embryo. Higher magnification view of quail mandibular primordia (dashed lines) in **(K)** whole-mount and **(L)** with RFP showing a size reduction on the treated (arrow) versus control side. **(M)** Quail embryo in whole mount at HH27 following unilateral *in ovo* injection of a control anti-GFP morpholino (MO) at HH18. At higher magnification in **(N)** whole-mount and **(O)** in schematic view, the mandibular primordia (dashed lines) appear the same size on the treated (striped) and control (green) sides. **(P)** Quail embryo in whole mount at HH27 following unilateral *in ovo* injection of an anti-*Gas1* morpholino (MO) at HH18. At higher magnification in **(Q)** whole-mount and **(R)** in schematic view, the mandibular primordia (dashed lines) appear smaller on the treated (striped) versus control (green) sides. Significance is shown for comparisons between control and treatment groups as denoted by brackets and p-values are as indicated. P-values ≥ 0.05 are considered not significant (ns). n ≥ 2 for each group and data point. Error bars represent SEM.

We also knocked-down *Gas1* expression in quail embryos by injecting an anti-*Gas1* morpholino into the right side of the mandibular primordia at HH18. We find that anti-GFP control morpholinos had no observable effect on jaw morphology (Figure 7M) as the left and rights sides appear equivalent (Figure 7N and 7O). However, in the anti-*Gas1* morpholino-injected embryos we observe left-right asymmetry in the developing jaw (Figure 7P, 7Q, and 7R). Taken together, our morphological, molecular, and cellular data indicate that changes in the response of NCM to SHH signaling and species-specific differences in *Gas1* expression may be a mechanism through which NCM establishes and modulates jaw size during development, disease, and evolution.

## Discussion

### Early developmental parameter values establish species-specific jaw size

In his highly influential book *On Growth and Form*, originally published in 1917, D’Arcy Thompson addressed the origins of changes in anatomical size and shape, and he emphasized the inseparable connection between morphology (i.e., form) and function (Thompson, 1917; Woronowicz and Schneider, 2019). Thompson argued that changes in coordinate scale could alter body plan proportions and act as a major driving force for evolutionary transformations (Thompson, 1917). Such allometry and its mathematical approach (i.e., power law equation) has been used to describe the relationship between measured quantities (e.g., morphogen diffusion rate, gene expression levels, rate of maturation, etc.), to model differential growth (i.e., changes in size and shape) of different parts of one organism, as well as to compare variation in the same or multiple body parts across species (Huxley, 1924; Huxley, 1932; Huxley and Teissier, 1936; Longo and Montévil, 2014; Schneider, 2018a).

Our previously published data together with our quantitative analysis of early embryonic stages in three birds with distinct jaw sizes (i.e., duck, chick, and quail) indicate that species-specific differences in the mandibular arch arise very early during development. We have shown that the migratory (i.e., *Sox10*-positive) population of NCM is about 25% larger in duck than in quail and that cell cycle length for mandibular mesenchyme is 11 hours in quail and 13.5 hours in duck (Fish et al., 2014). Thus, the initial size of the migratory NCM population that is allocated to the mandibular primordia, the amount of absolute time between developmental stages, and differences in cell cycle length can be major contributing factors to the establishment of species-specific jaw size (Fish et al., 2014). For this reason, we used a simple power function (*f(x)*) = *ax^b^* to model the relative population size of duck and quail throughout early development and we find that we can replicate mathematically what we observe *in vivo* (Figure 1B to 1D). These findings support the conclusion that the regulation of time (both in terms of embryonic stage and absolute), progenitor number, and cell cycle length during development are key parameters for scaling species-specific jaw size throughout evolution (Schneider, 2018a). Similarly, much theoretical and experimental work has also emphasized how varying the values for these types of parameters can directly affect diffusion, condensation, adhesion, differentiation, and other critical biochemical and cell-cell interactions that underlie morphogenesis (Oster and Alberch, 1982; Alberch, 1985; Alberch, 1989; Hall and Miyake, 1992; Hall and Miyake, 1995; Hall and Miyake, 2000).

### Transcriptional activation of the SHH pathway is species-specific and NCM-mediated

Our results reveal that the expression of members and targets of the SHH pathway is species-specific on multiple levels, starting with expression of the ligand, its receptors, as well as downstream effectors such as the *Gli* transcription factors. We observe distinct temporal patterns and changes in levels of expression for duck and quail. For example, *Shh, Ptch1, Gli1, Boc, Cdon, Smo, Gli2,* and *Gli3* have higher peak expression levels in duck relative to quail (Figure 2A, 2B, 2D, and Supplemental Figure 1). Interestingly, chick closely follows the gene expression patterns of quail from HH15 to HH18 and shifts to more duck-like expression from HH21 to HH27. Similarly, the ratios representing transcriptional activation of the SHH pathway suggest that chick is intermediate between duck and quail for *Ptch1*, but about the same as quail for *Gli1*. Further, analysis of our quail-duck chimeric data uncovers NCM-mediated gene expression at various hierarchical levels of the SHH pathway. These data show clear distinction between genes that are to various extents regulated by NCM (i.e., *Gas1*, *Gli1*, *Ptch1*, *Gli2*, and *Gli3*; Figure 3E, and 3G to 3J) and genes not affected by NCM (i.e., *Shh*, *Boc*, *Cdon*, and *Smo*; Figure 3B to 3D, and 3F). Such results may reflect the observation that the jaw primordia of chick fall somewhere between duck and quail in terms of absolute size, but the jaw primordia of chick and quail are more similar in shape (Smith et al., 2015). Accordingly, we can speculate that the axes of growth that generate species-specific size and shape may be differentially affected by *Ptch1-*versus *Gli*-mediated activation of the SHH pathway. Other studies have shown that various aspects of the SHH pathway can affect tissue size and shape such as changes in *Shh* enhancers (Kvon et al., 2016), variation in levels of SHH ligand, (Young et al., 2010; Xu et al., 2015), differential regulation of SHH receptors (Lopez-Rios et al., 2014; Xavier et al., 2016; Echevarría-Andino and Allen, 2020), transcriptional activity of target genes (Chang et al., 2016; Uygur et al., 2016), and variation in interacting co-factors (Elliott et al., 2020; Swartz et al., 2021).

Our analyses uncover species-specific variation in expression throughout early development and reveal that duck, chick, and quail have their own unique patterns at different hierarchical levels of the SHH pathway. Previously, SHH signaling activity has been shown to contribute to continuous variation in upper jaw morphology (Young et al., 2010; Swartz et al., 2012; Hu et al., 2015a; Hu et al., 2015b). In the mid and upper face, subtle differences in SHH dosage levels regulate facial width, where one extreme is facial clefting and the other is holoprosencephaly. SHH signaling activity in the face is regulated by many pathway members, including SHH receptors and GLI proteins (Seppala et al., 2007; Seppala et al., 2014; Chang et al., 2016; Xavier et al., 2016; Echevarría-Andino and Allen, 2020). Our data supports a similar model for the lower jaw where GAS1 mediates SHH signaling activity and small differences in *Gas1* expression in mandibular NCM contribute to species-specific jaw size. Together these data support the notion that SHH signaling can be fine-tuned by modular alterations to distinct genes or proteins within the pathway.

### Gas1 affects the species-specific development of the mandibular primordia

Our gene expression analyses as well as activation and inhibition experiments demonstrate that out of all the SHH pathway members that we examined, *Gas1* shows the greatest interspecific variation with the most distinct species-specific patterns detected during early developmental stages (Figure 2, Figure 4, and Figure 5; Supplemental Figure 1 and Supplemental Figure 2). Furthermore, our quail-duck chimeras reveal that *Gas1* expression is regulated by NCM (Figure 3E). In contrast, our *in situ* hybridization analyses show that the spatial domains of *Gas1* expression are similar in all three species at HH21 and HH27 (Figure 5A to 5F). Additionally, gene expression analyses of mesenchyme and epithelia in duck and chick at HH24 and HH27 show similar *Gas1* expression levels (i.e., high in mesenchyme and low in epithelium) (Figure 5G to 5H). Thus, while the spatial patterns of *Gas1* expression are conserved across species, variation in the levels of expression may contribute to species-specific differences in the growth of the mandibular primordia. Moreover, the species-specific levels in *Gas1* expression correlate with cell cycle length, suggesting there may be a mechanistic relationship between *Gas1* expression and cell cycle length in the mandibular primordia.

Duck have the highest levels of *Gas1* expression and the longest cell cycle compared to quail and chick (duck *Gas1* expression is up to 75 times higher than quail depending on stage). We observe the lowest levels of *Gas1* expression in chick and the average cell cycle length for chick has been estimated to be around 10 hours in early generations of cells and depending on the cell type (Morris et al., 1979; Smith and Schoenwolf, 1987; Primmett et al., 1989; Morris and Cowan, 1995; Venters et al., 2008). Thus, chick may have the shortest cell cycle length compared to quail and duck. Moreover, our data are consistent with reports from studies in other organisms where *Gas1* has been shown to act as a negative regulator that plays a critical role in growth suppression by reducing cell proliferation (Sacilotto et al., 2015; Sarkar et al., 2018). Taken together, our results support a model whereby species-specific sensitivity to SHH signaling and differential expression of *Gas1* affect cell cycle length and proliferation dynamics in NCM, which in turn produces distinct growth trajectories and ultimately contributes to size differences in the mandibular primordia in chick, quail, and duck.

Since its identification in 1988 (Schneider et al., 1988), *Gas1* has been shown to play an important role in a range of biological contexts. For example, Gas1 inhibits the transition from G_0_ to S phase, induces growth arrest through p53 (Del Sal et al., 1992; Del Sal et al., 1995), and functions as a tumor suppressor gene by regulating apoptosis (Zamorano et al., 2004). During embryogenesis, Gas1 acts as a positive regulator of cell proliferation and survival (Lee and Fan, 2001; Lee et al., 2001; Liu et al., 2001). Likewise, our overexpression experiments demonstrate that *Gas1* levels are critical for maintaining the amount of NCM and the size of the mandibular primordia. Knocking down *Gas1* expression also affects the size of the mandibular primordia as evidenced by the left-right asymmetry in embryos treated with morpholinos. Future research dedicated to elucidating how differential *Gas1* expression regulates cell number in the mandibular primordia and how *Gas1* modulates growth among different species has the potential to illuminate an important molecular mechanism regulating jaw size during development, disease, and evolution.

## Materials and Methods

### The Use of Avian Embryos

For all experiments, we adhered to accepted practices for the humane treatment of avian embryos as described in S3.4.4 of the AVMA Guidelines for the Euthanasia of Animals: 2013 Edition (Leary et al., 2013). Embryos were not allowed to hatch and no live vertebrate animals were used in this study. Fertilized eggs from white Pekin duck (*Anas platyrhynchos*), domestic chick (*Gallus gallus domesticus*), and Japanese quail (*Coturnix coturnix japonica*) were obtained from a commercial supplier (AA Labs, Westminster, CA, USA) and incubated at 37.8 °C and 85 to 87% humidity until reaching stages appropriate for surgery, manipulations, collection, and/or analysis.

To visualize embryos at early stages, a small amount of sterile 0.5% Neutral Red solution (N4638, Sigma-Aldrich, MilliporeSigma, St. Louis, MO, USA) was brushed lightly over the embryo with a blunt glass rod. Embryos were staged using the Hamburger and Hamilton (HH) staging system, a well-established standard that is based on external morphological characters and that is independent of body size and incubation time (Hamburger and Hamilton, 1951; Hamilton, 1965; Ricklefs and Starck, 1998; Starck and Ricklefs, 1998; Ainsworth et al., 2010). Absolute times of incubation to reach a particular embryonic stage for each species are in Figure 1A, 3A and Supplemental Table 1.

### Generation of Chimeric Embryos

Quail and duck were incubated until stage matched at HH9.5 (about 32 hours for quail and 55 hours for duck) (Figure 1A). To generate chimeric “quck” embryos, bilateral grafts of neural folds containing presumptive NCM from the anterior hindbrain and midbrain (dark green, Figure 3A) were excised from quail donors and transplanted into stage-matched duck hosts with a comparable region of tissue removed using flame-sharpened tungsten needles (Tungsten wire, 7190, A-M Systems, Sequim, WA, USA) and hand-made Spemann pipettes (Fish and Schneider, 2014a). Orthotopic control grafts were performed as done previously (Noden, 1983; Schneider, 1999; Schneider et al., 2001; Helms and Schneider, 2003; Lwigale and Schneider, 2008). After surgery, eggs were sealed with tape (3M Scotch Transparent Film Tape 600, 3M United States, St. Paul, MN, USA) and placed in the incubator until reaching stages appropriate for collection and analysis.

### Isolation of Mandibular Primordia

Using forceps, mandibular primordia were cut along the proximal junction at each side of the maxillary primordia and placed into RNase-free ice-cold 1x PBS (BP3991, Fisher Scientific, Hanover Park, IL, USA). For RNA extraction, isolated mandibular primordia were transferred into 1.5 mL microcentrifuge tubes with as little 1x phosphate buffered saline (PBS) as possible. Samples for RNA extraction were flash frozen on dry ice in 100 % ethanol and stored at -80 °C until ready to process.

### Quantification of Mandibular Mesenchyme

Mandibular mesenchyme was quantified in HH18, HH21, HH24, and HH27 duck, chick, and quail embryos. Trypsin-pancreatin solution was made at room temperature by dissolving (i.e., stirring for 15 minutes) 2.25 g of trypsin (T7409-100G, Lot # 029K7012, Sigma-Aldrich, MilliporeSigma, St. Louis, MO, USA) and 0.75 g of pancreatin (P1625-100G, Lot # SLBP2575V, Sigma-Aldrich, MilliporeSigma, St. Louis, MO, USA) in Hanks’ Balanced Salt Solution (HBSS; 14175095, ThermoFisher Scientific by Life Technologies Corporation, Grand Island, NY, USA). Pre-made trypsin-pancreatin solution was stored at -20 °C in 5 mL single use aliquots. The solution was thawed prior to dissection, filter sterilized using a 5 mL syringe (309646, BD Vacutainer Labware Medical, Fisher Scientific, Hanover Park, IL, USA) with 0.2 µm filter (431219, Corning Life Sciences Plastic, Lowell, MA, USA), and kept on ice until ready for use.

Isolated mandibular primordia were incubated in 0.5 mL trypsin-pancreatin solution at 4 °C on a rocker. Incubation length varied for each stage and species due to differences in size. Incubation times are listed in Supplemental Table 20. The trypsin-pancreatin solution was replaced with 1 mL of ice-cold HBSS to stop the enzymatic reaction. Samples were washed for 5 minutes at 4 °C on rocker. The solution was replaced with 1 mL of ice-cold Dulbecco’s Modified Eagle’s Medium (DMEM; 11965092, ThermoFisher Scientific by Life Technologies Corporation, Grand Island, NY, USA) and samples were kept on ice.

Mandibular epithelium was removed using two pairs of sharp forceps. Isolated mandibular mesenchyme was transferred into 1.5 mL microcentrifuge tubes containing 200 µL for HH18, 500 µL for HH21 and HH24, and 900 µL for HH27 of DMEM; homogenized by pipetting until all clumps were separated into single cells; and kept on ice. Mandibular mesenchyme (i.e., cell number per mandibular primordium) was counted using a hemacytometer (1492, Hausser Scientific Partnership, Fisher Scientific, Hanover Park, IL, USA) in combination with MIPAR software (Tyler and Hall, 1977; Sosa et al., 2014). Mean, standard error of the mean (SEM), number of biological replicates for each species, and data points are listed in Supplemental Table 2.

### Modelling Relative Population Size

To model cell counts per mandibular primordia, the size of *Sox10*-positive cell population, cell cycle length, and absolute developmental time were taken into account. The relative cell population size at HH13, HH15, HH18, HH21, HH24, and HH27 in duck and quail was calculated as *ax^b^*, where *x* represents cell division (i.e., doubling event when a mother cell divides in two daughter cells) and is equal to 2, *b* represents the number of cell cycles that the cell population for a particular species is able to go through within a period of time (e.g., from HH18 to HH21) and is calculated as follows 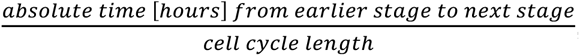, and *a* is the relative cell population at a previous developmental time point (i.e., earlier stage). The starting relative population size at HH13 was set to 1 for quail and calculated for duck by dividing the duck *Sox10*- positive cell population by quail *Sox10*-positive cell population using previously published values (Fish et al., 2014).

### Extraction of RNA

Total RNA was extracted from mandibula primordia at HH15, HH18, HH21, HH24, and HH27, mandibular mesenchyme, mandibular epithelium, explant tissue cultures, *Gas1* overexpression stable cell lines (DF-1), and whole heads and forelimbs at HH24 using the Arcturus PicoPure RNA Isolation kit (KIT0214, Applied Biosystems, Life Technologies Corporation, Grand Island, NY, USA) and following the manufacturer’s protocol with modifications as described in Supplemental Methods.

### Preparation of cDNA Libraries

Depending on the total RNA amount per sample available 100, 200, or 400 ng of total RNA was reverse transcribed to cDNA in a 20 µL reverse transcription reaction using 5 µL of iScript reverse transcription supermix (1708841, Bio-Rad Laboratories - Life Sciences, Hercules, CA, USA), a corresponding volume of total RNA template (calculated with the formula: 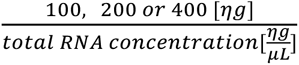), and sufficient nuclease-free water to bring the total volume up to 20 µL.

The cDNA synthesis reaction involved four steps: 25 °C for 5 minutes, 42 °C for 30 minutes, 85 °C for 5 minutes, and 4 °C hold in a thermocycler (model C1000 Touch Thermal Cycler, Bio-Rad Laboratories - Life Sciences, Hercules, CA, USA). Lid temperature was set to 105 °C. cDNA libraries were stored at -20 °C.

### Designing and Validating Primers for qRT-PCR

Species-specific primers were designed for *Shh*, *Ptc1*, *Cdon*, *Boc*, *Smo*, *Gas1*, *Gli1*, *Gli2*, and *Gli3* using Geneious Prime (Biomatters, San Francisco, CA, USA) (i.e., bioinformatics suite that incorporates Primer 3 (Koressaar and Remm, 2007; Untergasser et al., 2012)). Primer sequences are in Supplemental Table 21. Design criteria for primers included a product size between 70 and 200 bp (with optimal being 150 bp); a primer size minimum of 18 bp and maximum of 27 bp (with optimal being 20 bp); a Tm minimum of 50 °C and maximum of 65 °C (with optimal being 60 °C); a GC content minimum of 45% and maximum of 60% (with optimal being 50%); a maximum Tm difference for primer pair of 5 °C; a maximum dimer Tm of 40 °C; a maximum 3’ stability of 9; a GC clamp of 1; and a maximum Poly-X of 3. The SantaLucia 1988 formula and salt concentration were set for Tm calculation with concentration settings of 50 mM for monovalent cations, 3 mM for divalent cations, 500 nM for oligos, and 0.8 mM for dNTPs. Gene sequences were checked for SNPs and those regions were avoided. Primer pairs for all genes spanned the same region for all three species. For genes with multiple splice variants, primers were designed to regions present in all known isoforms. To prevent protentional amplification of genomic DNA, all primer pairs, with the exception of *Gas1* (which has a single exon), were designed to span exon-exon junctions. All potential primer pairs were blasted using Primer Blast (Ye et al., 2012) against the genomes of duck, chick, and quail to avoid cross reactivity and amplification of unintended target sequences. Selected primer pairs were ordered from IDT (Integrated DNA Technologies, Coralville, IA, USA) with standard desalting. Lyophilized primers were diluted in an appropriate amount of RNase/DNase free water to make a stock concentration of 100 µM, which was stored at -20 °C. Prior to use, the stock solution was diluted 1:10 (i.e., working concentration).

To validate species-specific primers, RNA was extracted from whole heads and forelimbs of HH24 duck, chick, and quail embryos as described above. 400 ng of RNA was reverse transcribed to cDNA and cDNA was diluted to a concentration of 2 ng/µL (1x). Standard curves with five serial dilutions (1:4, 1:16, 1:64, and 1:256) for each species-specific primer pair were generated from a 25 µL reaction mixture (1.5 µL of forward primer at working concentration, 1.5 µL of reverse primer at working concentration, 4 µL of cDNA, 6 µL of RNase/DNase free water, and 12.5 µL of iQ SYBR-Green Supermix (1708884BUN, Bio-Rad Laboratories - Life Sciences, Hercules, CA, USA)), run in technical triplicates on hard-shell PCR white 96-well plates (HSP9601, Bio-Rad Laboratories - Life Sciences, Hercules, CA, USA), sealed with optically clear microseal film for PCR plates (MSB1001, Bio-Rad Laboratories - Life Sciences, Hercules, CA, USA), and following the protocol described below. Controls for each run included no reverse transcription and no cDNA. Melt curves were checked for primer specificity (i.e., single product) and for excluding samples with potential genomic DNA contamination. To confirm the correct products size, 1x samples were run on a 1 % Agarose gel (BP1356-500, Fisher Bioreagent, Fisher Scientific, Hanover Park, IL, USA) in 1x TAE buffer at 110 V for 30 minutes. The Quick-Load 100bp Ladder (N0467S, New England BioLabs, Ipswich, MA, USA) was used as a size reference.

For selected primers, products were amplified in a 25 µL PCR reaction mixture using a thermocycler (model C1000 Touch Thermal Cycler, Bio-Rad Laboratories - Life Sciences, Hercules, CA, USA). PCR reaction reagents included 2.5 µL of 10x Buffer (42-800B3, Apex Bioresearch Products, Genesee Scientific, El Cajon, CA, USA), 0.5 µL of dNTPs (17-106, PCR Biosystems, Genesee Scientific, El Cajon, CA, USA), 0.75 µL of MgCl_2_ (42-800B3, Apex Bioresearch Products, Genesee Scientific, El Cajon, CA, USA), 1.25 µL of DMSO (D128-500, Fisher Chemical, Fisher Scientific, Hanover Park, IL, USA), 0.125 µL of Taq (42-800B3, Apex Bioresearch Products, Genesee Scientific, El Cajon, CA, USA), 1.25 µL of forward primer at working concentration, 1.25 µL of reverse primer at working concentration, 13.0 µL of DNase/RNase-free water, and 4.375 µL of cDNA at the concentration of 2 ng/µL. The protocol for product amplification was: Step 1, 94 °C for 2 minutes; step 2, 94 °C for 2 minutes; step 3, 57.5 °C for 30 seconds; step 4, 72 °C for 1 minute; steps 2 to 4 were repeated 39 times; step 5, hold at 4 °C. Lid temperature was set to 105 °C. To verify primer pair products, PCR product clean up and sequencing was done by Molecular Cloning Laboratories (www.mclab.com, South San Francisco, CA, USA).

### Analysis of Gene Expression by qRT-PCR

4.6 µL aliquots of cDNA (1 ng/µL) were prepared and stored in 0.2 mL snap strip PCR tubes with dome caps (490003-692, GeneMate – BioExpress, VWR International, Brisbane, CA, USA) at -20 °C prior to qRT-PCR plates preparation. Only a portion (i.e., volume necessary to run qRT-PCR for selected genes and primer pairs) of the total volume of working solution for each sample was aliquoted and the remaining volume stored in 0.5- or 1.5-mL microcentrifuge tubes at -20 °C until ready for the next usage to prevent the number of freeze-thaw cycles.

qRT-PCR was performed in a C1000 Touch Thermal Cycler with a CFX96 Real-Time System (1855196, Bio-Rad Laboratories - Life Sciences, Hercules, CA, USA). The reaction mixture for each well contained 1.25 µL of forward primer at working concentration, 1.25 µL of reverse primer at working concentration, 2 µg of cDNA (i.e., 2 µL at the concentration of 1 ng/µL), 10 µL of iQ SYBR-Green Supermix (1708884BUN, Bio-Rad Laboratories - Life Sciences, Hercules, CA, USA), containing dNTPs, iTaq DNA polymerase, MgCl_2_, SYBR Green I, enhancers, stabilizers, and fluorescein, and 5 µL of RNase-free water. Samples were run in technical duplicates on hard-shell PCR white 96- well plates (HSP9601, Bio-Rad Laboratories - Life Sciences, Hercules, CA, USA) sealed with optically clear microseal film for PCR plates (MSB1001, Bio-Rad Laboratories - Life Sciences, Hercules, CA, USA) using tested protocol. The protocol steps were following: Step 1, 95 °C for 3 minutes; step 2, 95 °C for 10 seconds; step 3, 60 °C for 30 seconds and a plate read; steps 2 and 3 were repeated 39 times; step 4, 95 °C for 10 seconds; step 5, melt curve of 60-90 °C for 5 seconds at each 0.5 °C with a plate read. Lid temperature was set to 105 °C. Melt curves were used to assess the specificity of primers and to exclude samples with potential genomic DNA contamination. Gene expression levels were quantified using the Pfaffl method (Pfaffl, 2001). Levels were normalized to *r18s*.

### Culture of Mandibular Primordia and Biochemical Treatments

For explant cultures, dissected mandibular primordia were processed as described previously (Eames and Schneider, 2008; Merrill et al., 2008). Briefly, isolated mandibular primordia were placed onto a small circular piece of nitrocellulose membrane filter (AAWP04700, MilliporeSigma, St. Louis, MO, USA), which was then transferred to a 50 mm Petri dish containing DMEM (11965092, ThermoFisher Scientific by Life Technologies Corporation, Grand Island, NY, USA). Nitrocellulose membrane filters were cut prior to use with a sterilized metal 6 mm pore diameter punch pliers. Mandibular primordia were positioned into the center of the membrane filter, which was then placed into a transwell plate (3413, Corning Life Sciences Plastic, Lowell, MA, USA) containing 325 µL of complete media (75% DMEM, 10% horse serum (16050122, ThermoFisher Scientific by Life Technologies Corporation, Grand Island, NY, USA), and 15% chick embryo extract ultrafiltrate (C3999, US Biological Life Sciences, Salem, MA, USA)) and incubated at 37 °C with 5% CO_2_.

To make SHH stock solution lyophilized human recombinant (r) SHH protein (SHH-005/SHH-100, Lot #1362 for mandibular primordia explants and Lot #1371 for cell culture, StemRD, Burlingame, CA, USA) was reconstituted in sterile water to a concentration of 100 ng/µL. Stock solution was stored at -80 °C in 7.15 µL aliquots. 45 µL aliquots of rSHH protein was diluted in complete media to achieve final concentrations of 0.1, 1, 10, 100, or 1000 ng/mL. To make cyclopamine stock solution, lyophilized Cyclopamine-KAAD (Lot # 2944661, 239804, MilliporeSigma, Burlington, MA, USA) was reconstituted following the manufacturer’s instructions to a concentration of 4 ng/µL and stored at -20 °C in 20 µL aliquots.

Mandibular primordia were placed with the oral side down on the membrane for control and rSHH treatments, and with the aboral side on the membrane for cyclopamine treatments. 45 µL of rSHH protein at each concentration were added to the side of each well. rSHH protein was mixed into the complete media by rotating the 24-well plate. For cyclopamine treatment, 40 µL of complete media was added to the side of each well and 5 µL of the cyclopamine stock solution was pipetted on top of the mandibular primordia. For control samples, 45 µL complete media was added to the side of each well. The final volume per well, after adding rSHH protein at each concentration, cyclopamine, or complete media was 370 µL.

After 24 hours, explants were removed from the transwells and transferred into 50 mm Petri dishes containing RNase-free ice-cold 1x PBS. Explants were carefully detached from the cellulose membrane using a pair of forceps and a 1 mL syringe (329654, BD Vacutainer Labware Medical, Fisher Scientific, Hanover Park, IL, USA) with a 30 gauge by 0.5-inch needle (305106, BD Vacutainer Labware Medical, Fisher Scientific, Hanover Park, IL, USA). Each explant was transferred into a 1.5 mL microcentrifuge tube with snap cap with as little 1x PBS as possible, flash frozen on dry ice, and stored at -80 °C. Harvested explants were processed for qRT-PCR as described above.

### Generating Probes for In Situ Hybridization

PCR primers were designed against the coding region of *Gas1* using Primer3 (Untergasser et al., 2012) targeting a conserved region in duck, chick, and quail that would generate a ∼650 bp probe. Primers were synthesized with and without T7 RNA polymerase promoter consensus sequence 5’-TAA TAC GAC TCA CTA TAG GG -3’ as a primer tail. The forward primer with the T7 RNA polymerase promoter was used to generate sense template and the reverse primer with T7 RNA polymerase promoter was used to generate antisense template. Q5 Hot Start High-Fidelity DNA Polymerase (M0493L, New England BioLabs, Ipswich, MA, USA) was used to amplify the templates for *in vitro* transcription. *In vitro* transcription was carried out using T7 RNA polymerase (M0251S, New England BioLabs, Ipswich, MA, USA) following the manufacturer’s directions with digoxigenin (DIG) RNA Labeling Mix (11277073910, Roche, Basel, Switzerland) to generate DIG labelled probe. RNaseOUT (10777019, Invitrogen, Carlsbad, CA, USA) was used to protect the RNA from degradation during synthesis. *In vitro* transcription was carried out at 37 °C for 2 hours. Following *in vitro* transcription, DNA was degraded using DNase I (4716728001, Roche, Basel, Switzerland). RNA probes were then precipitated with LiCl and ethanol. RNA probes were resuspended in RNAse free MilliQ water and then mixed with an equal volume of formamide (VWRV0606-100ML, VWR, Radnor, PA, USA) to stabilize the RNA. Probes were stored at -20 °C until ready for use. Primer sequences for species-specific *Gas1* probes are listed in Supplemental Table 22.

### Analysis of Gas1 Expression by In Situ Hybridization

Mandibular primordia were isolated from duck, chick, and quail embryos at HH21 and HH27 using forceps and placed in RNase-free ice-cold 1x PBS. Samples were transferred into 15 mL conical tubes and fixed in 4% paraformaldehyde (PFA) at 4 °C for two hours. Samples were washed three times for 30 minutes at room temperature in 1x PBS followed by 30 minutes washes in 25%, 50%, and 75% methanol in 1x PBS and three times in 100% methanol (A412-4, Fisher Chemical, Fisher Scientific, Hanover Park, IL, USA). Samples were stored at -20 °C in 100% methanol until processing.

Whole mount *in situ* hybridization was performed following published protocols (Merrill et al., 2008; Fish et al., 2014). A solution of Proteinase K (P6556, Sigma-Aldrich, MilliporeSigma, St. Louis, MO, USA) in 1x PBS was used at a concentration of 10 µg/mL for HH21 embryos and 20 µg/mL for HH27 embryos. Incubation times were 25 minutes for HH21 embryos and 15 minutes for HH27 embryos. Following the color reaction, developed for 2 hours at room temperature then for 14 hours at 4 °C, mandibular primordia were imaged on a stereo dissecting microscope under brightfield illumination (model MZFLIII-TS, Leica Microsystems, Buffalo Grove, IL, USA).

### Cloning Full-Length Gas1

Full-length cDNA synthesis from RNA was performed using Maxima H Minus reverse transcriptase (K1651, Thermo Scientific, Fisher Scientific, Hanover Park, IL, USA) following the manufacturer’s directions with 2 µg of total RNA and 100 pmol of d(T)20 VN primer. The cDNA synthesis reaction was carried out at 50 °C for 30 minutes, 55 °C for 10 minutes, 60 °C for 10 minutes, 65 °C for 10 minutes, and 85 °C for 5 minutes. Full length duck, chick, and quail *Gas1* was amplified by PCR in a thermocycler (model 2720 Thermal Cycler Applied Biosystems, Carldbad, CA, USA) using Q5 Hot Start High-Fidelity DNA Polymerase (M0493L, New England BioLabs, Ipswich, MA, USA) and cloned using CloneJET PCR Cloning Kit (K1231, Thermo Scientific, Fisher Scientific, Hanover Park, IL, USA). Full-length duck, chick, and quail *Gas1* were confirmed by Sanger sequencing. *Gas1* was cloned into the pPIDNB plasmid, which integrates into the genome and is doxycycline (dox)-inducible (Chu et al., 2020), using AflII (R0520S, New England BioLabs, Ipswich, MA, USA), PstI (R3140S, New England BioLabs, Ipswich, MA, USA), and NEBuilder HiFi DNA Assembly Master Mix (E2621L, New England BioLabs, Ipswich, MA, USA). All constructs were verified by Sanger sequencing and midiprepped for electroporation and/or transfection using PureLink Fast Low-Endotoxin Midi Kit (A36227, Invitrogen, Fisher Scientific, Hanover Park, IL, USA) (Chu et al., 2020). Empty pPIDNB plasmid was used as a control.

### Preparation of Micropipettes

Micropipettes for DNA injection were generated using a micropipette puller (model P-87 Flaming/Brown, Sutter Instrument Co., Novato, CA, USA). Borosilicate capillary glass without a filament and with an outside diameter of 1 mm and an inner diameter of 0.75 mm (B100 – 75 – 10, Sutter Instrument Co., Novato, CA, USA) was used. Program settings were as follows: Heat = 693, Velocity = 50, Pull = 100, Time = 250, Press = 300.

### The Use of Cell Culture and Generation of Stable Gas1-Expressing Lines

A fibroblast cell line (UMNSAH/DF-1) from embryonic chick (*Gallus gallus*) was obtained from the American Type Culture Collection (CRL-12203, ATCC, Manassas, VA, USA) and maintained as directed. DF-1 cells were cultured in complete media (i.e., DMEM (10-013-CV, Corning Mediatech Inc., Lowell, MA, USA) supplemented with 10 % Fetal Bovine Serum (FBS) (Lot # 283K18, 97068-085, VWR International, Brisbane, CA, USA) and 1x penicillin-streptomycin (15140122, ThermoFisher Scientific by Life Technologies Corporation, Grand Island, NY, USA)) at 37 °C with 5 % CO_2_. Cells were passaged twice a week.

DF-1 cells were transfected with lipofectamine 3000 reagent (L3000008, Invitrogen by Life Technologies Corporation, Grand Island, NY, USA) according to the manufacturer’s protocol. Transfections for integrating empty pPIDNB, pPIDNB-chick-*Gas1*, or pPIDNB-quail-*Gas1* were carried out in 6-well plates (353046, Corning Life Sciences, Corning, NY, USA) in technical duplicates. 500,000 cells were seeded per well 24 hours prior to transfection. Transfections were done using 2.5 µg of plasmid, 2.5 µg of pNano-hyPBase, and 10 µL of P3000 (Chu et al., 2020). Cells were incubated for 12 to 15 hours and then washed with 2 mL of a 0.25 % trypsin solution (25200056, Gibco, Fisher Scientific, Hanover Park, IL, USA) in 1X EDTA. Transfection efficiency was confirmed by visualizing fluorescence of constitutively active mNeonGreen in the pPIDNB plasmid (Chu et al., 2020) on a Nikon AZ100 C2+ Macro Confocal Microscope (Nikon Instrument, Inc., Melville, NY). Transfected cells from 2 wells (technical duplicates) were seeded into 250 mL cell culture flasks with vented caps (10062-860, VWR International, Brisbane, CA, USA).

Eight days post transfection, complete media was removed, cells were washed with 2 mL of trypsin and incubated in 3 mL of trypsin at room temperature until they started to detach from the bottom of the flask. Trypsin activity was inhibited by adding 5 mL of DMEM with 10% FBS. Cells were pipetted and passed through a 70 µm filter (352235, Corning Life Sciences, Corning, NY, USA). Cells were sorted on a FACSAriaII Flow Cytometer (BD Bioscience, San Jose, CA). All debris and dead cells were eliminated using FSC-A and SSC-A gating, doublets were excluded via gating discrimination using FSC-H and FSC-W, and two cell populations per construct (GFP^+^ and GFP^-^) were collected. Voltages used were 181 for FSC, 355 for SSC, and 363 for 530/30 blue C laser. Each cell population was collected into a 15 mL conical tube containing 3 mL of complete media. Three stable cell lines were generated: DF-1 containing an empty pPIDNB expression vector, DF-1 containing the chick *Gas1* pPIDNB expression vector, and DF-1 containing the quail *Gas1* pPIDNB expression vector. Each cell line was split into four flasks and incubated until reaching 90% confluency. Then cells from 3 flasks were detached as described, transferred into 15 mL conical tubes, and spun down at 200 x g for 5 minutes. Supernatant was carefully removed and cells resuspended in 3 mL of complete media. 1 mL of resuspended cells was transferred into a 2 mL cryogenic vial (430659, Corning Life Sciences Plastic, Lowell, MA, USA) and frozen overnight at -80 °C in a NALGENE Cryo 1 °C freezing container (5100-0001, ThermoFisher Scientific, Hanover Park, IL, USA) filled with isopropanol (423830025, Acros Organics, Fisher Scientific, Hanover Park, IL, USA). Cells were then stored at -140 °C in liquid nitrogen (CryoPlus Storage Systems model 7402, ThermoFisher Scientific, Hanover Park, IL, USA).

To quantify the number of cells per well, DF-1 cells were dissociated and counted using a hemacytometer (1492, Hausser Scientific Partnership, Fisher Scientific, Hanover Park, IL, USA) (Tyler and Hall, 1977). For quantification, DF-1 cells were imaged on a Zeiss Axiovert 40 CLF trinocular inverted microscope (Carl Zeiss AG, Göttingen, Germany) using fluorescence and phase contrast. All cell culture experiments were carried out using the same lot of DF-1 cells.

### Electroporation

A DNA solution containing 3 µg/µL pPIDNB (empty or containing *Gas1*), 1 µg/µL pNano-hyPBase plasmid, and 0.5 µL of 1% Fast Green was loaded into a glass micropipette (described above) and delivered with a picospritzer fluid injector (model PV830 Pneumatic PicoPump, SYS-PV830, World Precision Instruments, Sarasota, FL, USA). To target the presumptive NCM that migrates into the mandibular primordia (Fish et al., 2014), DNA solution was injected into the lumen of the dorsal neural folds from the anterior hindbrain to the anterior midbrain of HH8.5 duck and quail embryos. Homemade platinum electrodes (78-0085, Strem Chemicals Inc., Fisher Scientific, Hanover Park, IL, USA) mounted in an Adjustatrode holder (01-925-09, Intracel by Abbotsbury Engineering Ltd., St Ives, UK) were positioned on each side of the area pellucida, centered on the midbrain-hindbrain boundary. The distance between electrodes was set to 5 mm. The electrodes were overlayed with albumin to prevent drying and to facilitate conductivity.

*In ovo* electroporation was performed using a BEX CUY21EDITII Pulse Generator (CUY21EDIT2, BEX CO., LTD, Tokyo, Japan). Unilateral electroporations involved three square pulses delivered at 50 V with 10% decay for 1 ms spaced 50 ms apart followed by five square pluses at 10 V with 20% decay for 50 ms spaced 50 ms apart. Bilateral electroporations involved three square pulses at 50 V with 10% decay for 1 ms spaced 50 ms apart, three square pulses at 50 V with 10% decay for 1 ms spaced 50 ms apart in the reverse polarity, three five square pluses at 10 V with 20% decay for 50 ms spaced 50 ms apart followed by five square pluses at 10 V with 20% decay for 50 ms spaced 50 ms apart in the reverse polarity.

After electroporation, a small amount of albumin was added on top of the embryo to prevent desiccation. Eggs were sealed with tape and incubated at 38.3 °C. After 24 hours, electroporation efficiency was confirmed by visualizing fluorescence *in ovo* of constitutively active mNeonGreen in the pPIDNB plasmid (Chu et al., 2020) on a Nikon AZ100 C2+ Macro Confocal Microscope (Nikon Instrument, Inc., Melville, NY). Eggs were re-sealed with tape, incubated until reaching HH15, and treated with dox (see below).

### Treatment with Doxycycline

Stock solution of doxycycline hyclate (dox) (446060250, Acros Organics, Fisher Scientific, Hanover Park, IL, USA) was made up to final concentration of 1 mg/mL in filter sterilized water. Single use 200 µL aliquots of stock solution were stored at -20 °C.

For cell culture, dox stock solution was diluted to final concentrations of 0.1, 1, 5, 10, or 50 ng/mL in complete media (i.e., DMEM supplemented with 10% FBS and 1x penicillin-streptomycin). Dox treatment was performed by replacing complete media with prepared complete media containing appropriate dox concentrations. For control samples, complete media was replaced with fresh complete media without dox.

For *in ovo* dox treatments, egg volumes were considered to be 75 mL for duck and 8 mL for quail. A dox working solution was made up by mixing 7.5 µL for duck and 0.8 µL for quail of stock (i.e., 1mg/ml) dox solution with 750 µl for duck and 200 µL for quail of HBSS. Prepared working solution was gently pipetted through the egg window onto the vitelline membrane adjacent to the embryo and allowed to diffuse. The final dox concentration was 100 ng/mL. Eggs were sealed with tape and incubated until collection at HH18, HH21, HH24, and HH27. Duck mandibular primordia were isolated and imaged on a stereo dissecting microscope (MZFLIII-TS, Leica Microsystems, Buffalo Grove, IL, USA) under brightfield illumination or epifluorescent illumination to assess the distribution of *Gas1* overexpression as indicated by mScarlet fluorescence. The heads of quail at HH27 were stained with Hoechst 33342 Dye (see below) and imaged on a stereo dissection microscope under epifluorescent illumination (MZFLIII-TS, Leica Microsystems, Buffalo Grove, IL, USA).

### Injections with Vivo-Morpholinos

*In ovo* Vivo-Morpholino injections were performed using a 0.5 mM solution of Vivo-Morpholino (quail anti-*Gas1* or control anti-*GFP*) (Gene Tools, LLC, Philomath, OR, USA) in HBSS containing 0.01 % Phenol Red (417240050, Acros Organics, Fisher Scientific, Hanover Park, IL, USA). Vivo-Morpholino sequences are listed in Supplemental Table 23.

Vivo-Morpholino solution was injected into the right side of quail mandibular primordia at HH18 with pulled glass micropipettes loaded into a PicoNozzle (5430-10) and a picospritzer fluid injector (PV830, World Precision Instruments, Sarasota, FL, USA). After injection, eggs were sealed with tape and incubated until reaching HH27. The heads of treated embryos were stained with Hoechst 33342 Dye and imaged on a stereo dissection microscope under epifluorescent illumination (MZFLIII-TS, Leica Microsystems, Buffalo Grove, IL, USA).

### Staining with Hoechst Dye

To enhance contrast for phenotypic analysis, whole mount duck and quail embryos were stained with Hoechst 33342 Dye (62249, Lot# RF22228410, ThermoFisher Scientific, Hanover Park, IL, USA). Briefly, a 20 mM stock solution was diluted 1:1000 in 1x PBS to a working solution of 1 µg/mL. Samples were incubated in working solution for 48 hours at 4 °C on a rocker.

### Capture and Adjustment of Images

Brightfield and epifluorescent images were acquired with SPOT Insight 2.0 Firewire Color Mosaic camera (IN1820, Model 18.2.x) and SPOT image capture software (SPOT Imag3ing, Diagnostic Instrument, Inc., Sterling Heights, MI, USA). Multiple image planes were combined in Helicon Focus (version 7.6.1 Pro, Helicon Soft Ltd., 2000, Kharkiv, Ukraine). Images were adjusted in Adobe Photoshop 2020 (version 21.2.2) to normalize for exposure, brightness, contrast, saturation, and color balance across samples. Figures were assembled in Adobe Illustrator 2020 (Version 24.2.3). To label data points, the following colors were used: duck = violet/Medium Orchid, # D344DD; chick = yellow/Golden Dream, #F2D33C; quail = green/Magic Mint, #9FF4BA; and quck = blue/Calypso, #3C6B87.

### Methods for Determining Statistical Significance

Prism (v.9.1.0) software (GraphPad, San Diego, CA, USA) was used to perform statistical tests and determine significance. Unpaired multiple t-tests were performed at each time point for analysis of mandibular mesenchyme population size, gene expression levels *in vivo*, as well as gene expression ratios. The Holm-Šídák method was used to correct for multiple comparisons. Ordinary one-way ANOVA was performed for analysis of gene expression and gene expression ratios between duck, quail, and quck at quck embryonic stage HH21 and HH24, for analysis of gene expression following treatments in mandibular primordia explants and DF-1 cells, as well as for quantification of DF-1 cells when comparing different treatments to control samples. The Dunnett’s multiple comparison test was used to correct for multiple comparisons.

Error bars denote SEM. Multiplicity adjusted p-values were used to determine significance, and p values are indicated on the data points and/or legends of each figure. Formal power analyses were not conducted. The number of biological replicates for each data point and/or treatment group was between 2 and 17, specific number of biological replicates per data point and p-values are listed in the Supplemental Tables 2, 3, and 6 to 20.

## Acknowledgements

We thank Sophie E. Creuzet, Katherine C. Woronowicz, Tamara Alliston, Ralph S. Marcucio, Spenser S. Smith, Jessye Aggleton, and Evelyn E. Schwager for technical support and comments on the manuscript; Angeline K Chemel, Hugo Sugier, and Nicolas Rose for laboratory assistance; Thomas Dam at AA Lab Eggs for fertilized duck, chick, and quail eggs; the UCSF Biological Imaging Development CoLab (BIDC) for microscopy and data analysis support; and the UCSF Parnassus Flow Cytometry Core (PFCC, RRID:SCR_018206) for assistance generating Flow/Mass Cytometry data.

## Author contributions

Z.V., J.L.F, and R.A.S. conceived the project; Z.V., D.B.C., A.N., and J.L.F. performed the experiments; Z.V., A.N., and J.L.F. compiled the data; Z.V. and R.A.S. designed the experiments; Z.V. analyzed the data; Z.V. and R.A.S. co-wrote the manuscript and all authors edited and approved the final draft.

## Data Availability

Datasets supporting the results of this study that are not presented in the Supplemental Tables are available upon request from the corresponding author (R.A.S.). The GenBank accession number for *Gas1* nucleotide sequence is MW036691. The stable and inducible overexpression plasmids used in this study are available for distribution on Addgene (https://www.addgene.org/browse/article/28216052) subject to the terms of the original licenses under which they were obtained.

## Supplemental Methods

### RNA extraction

Steps for total RNA extraction using Arcturus PicoPure RNA Isolation kit (KIT0214, Life Technologies Corporation, Grand Island, NY, USA) were as follows:

1. RNA extraction

a. Whole mandibular primordia and explant tissue culture: 85 µL of extraction buffer was added to each sample.
b. Cell culture: Complete media was removed from wells and cells were washed twice with 300 µL of RNase-free 1x PBS. 150 µL of extraction buffer was added per well. Cells were scraped off of the well with 200 µL pipette and transferred into 1.5 mL microcentrifuge tube. Sample homogenization was carried out in a Bead Mill 24 Homogenizer (15-340-163, Fisher Scientific, Hanover Park, IL, USA) at 4 m/s for 15 s (C = 1, D = 0). Samples were spun down for 2 minutes at 16,000 RCF then incubated in water bath at 42 °C for 30 minutes. Samples were vortexed for 2 s in the middle of the incubation.
2. RNA isolation Extraction tubes were pre-treated following manufacturer’s directions. 85 µL of RNase-free 70 % ethanol, made by diluting 100 % ethanol (BP28184, Fisher Scientific, Hanover Park, IL, USA), was added to the tissue/cell extract and mixed by pipetting 20 times up and down. The extract was transferred into pre-conditioned purification column and centrifuged following manufacturer’s directions. 180 µL of RNase-free 70 % ethanol was added onto the column and centrifuged for 1 minute at 16,000 RCF. The rest of the protocol was carried out following manufacturer’s directions. Residual genomic DNA was removed in on-column step using RNase-free DNase Set (79254, Qiagen, Germantown, MD, USA), 5 µL of DNase I mixed with 35 µL of RDD buffer. Total RNA was eluted in 21 µL of Elution buffer for whole mandibular primordia at HH18, 21, 24, and 27 and explant tissue culture; and in 16 µL of Elution buffer for whole mandibular primordia at HH15 and cell cultures.

Concentration and purity of RNA were assessed using a NanoDrop spectrophotometer (model ND-1000, ThermoFisher Scientific by Life Technologies Corporation, Grand Island, NY, USA) and/or Qubit 4 fluorometer (Q33222, Invitrogen, Hanover Park, IL, USA) according to the manufacturer’s instructions. Total RNA extracted in Elution buffer was stored at -80 °C.

---

**Supplemental Figure 1.**
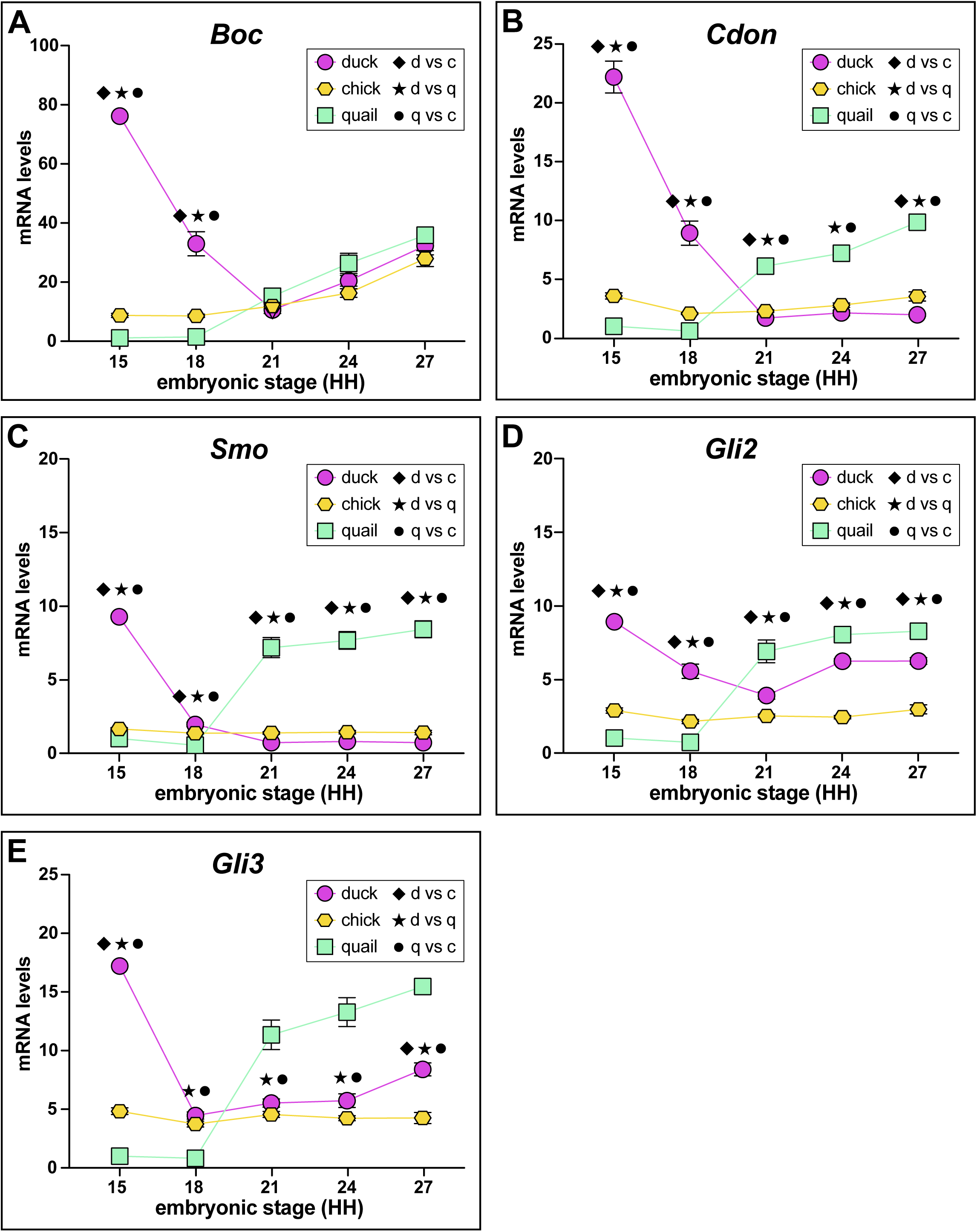
Expression of SHH pathway members in mandibular primordia of duck, chick, quail, and chimeric quck. Relative mRNA levels of **(A)** *Boc*, **(B)** *Cdon*, **(C)** *Gli2*, **(D)** *Gli3*, and **(E)** *Smo* in mandibular primordia of duck (violet), chick (yellow), and quail (green) at HH15, HH18, HH21, HH24, and HH27. Expression levels were assayed by qRT-PCR and normalized to *r18s*. Significance is shown (p-value < 0.02, n ≥ 6 for each group and data point, and error bars represent SEM) for comparisons between different species at the same embryonic stage (i.e., diamond symbol for duck versus chick, asterisk symbol for duck versus quail, and full circle symbol for quail versus chick).

**Supplemental Figure 2.**
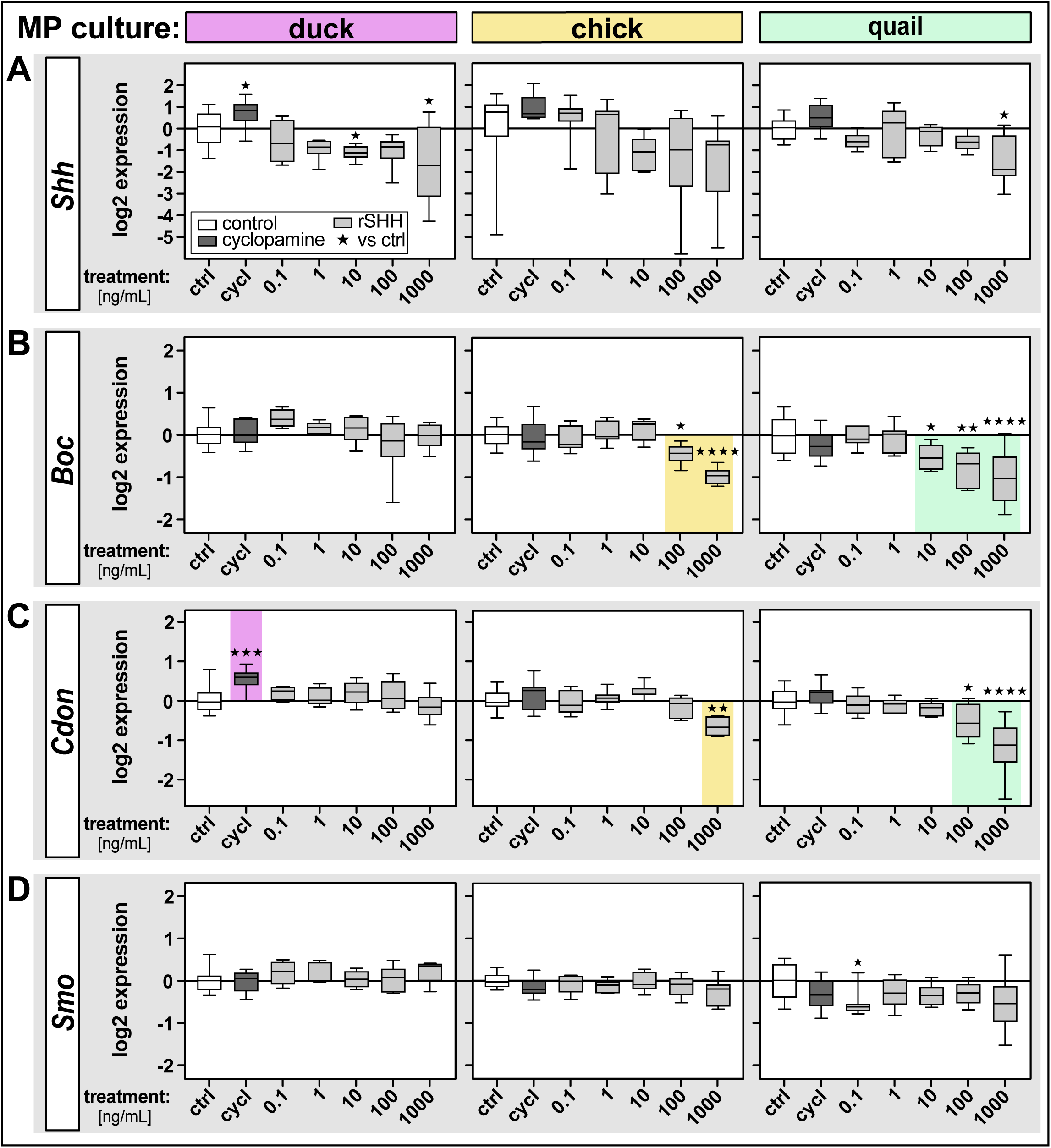
Effects of inhibition and activation of the SHH pathway in mandibular primordia of duck, chick, and quail. Mandibular primordia from duck (violet), chick (yellow), and quail (green) embryos were harvested at HH21, placed in culture, and treated with cyclopamine (cycl, dark grey) to inhibit the SHH pathway or with five serial dilutions of recombinant (r) SHH protein (light grey) to activate the SHH pathway. Box plots show relative levels of mRNA expression (on the y-axis in log_2_ scale) for SHH pathway members including **(A)** *Shh*, **(B)** *Boc*, **(C)** *Cdon*, and **(D)** *Smo* 24 hours after treatment. Control (ctrl, white) and treatment groups are shown on the x-axis. Expression levels were assayed by qRT-PCR and normalized to *r18s*. Significance is shown for comparisons between control and treatment groups (i.e., cyclopamine or rSHH) within the same species as indicated by colored shading and by the following symbols for p-values: * p < 0.05, ** p < 0.01, *** p < 0.001, and **** p < 0.0001. P-values ≥ 0.05 are considered not significant. n ≥ 4 for each group and data point.

**Supplemental Figure 3.**
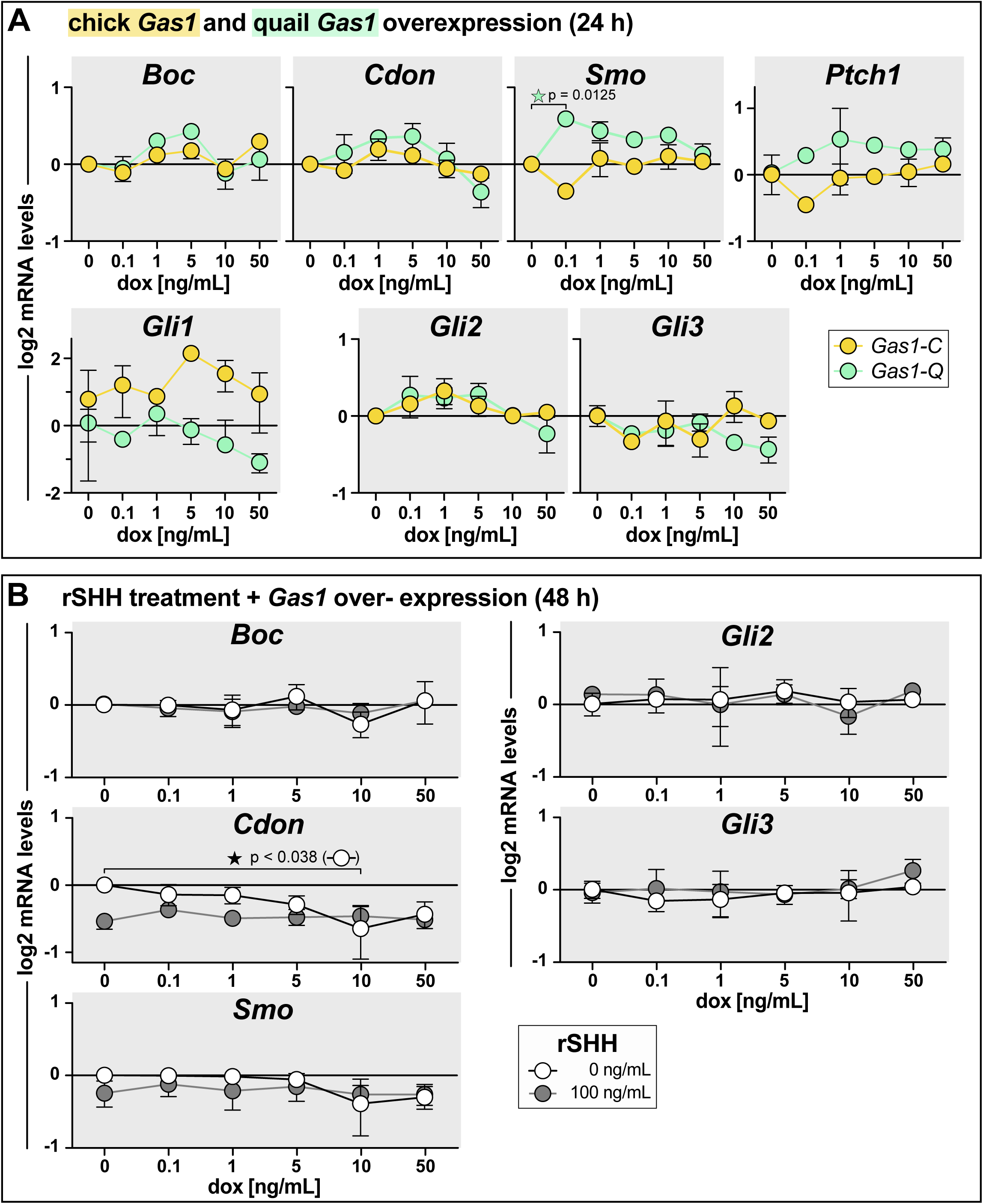
Effects of SHH pathway activation and *Gas1* overexpression in cell culture. **(A)** Relative levels of mRNA expression (on the y-axis in log_2_ scale) for *Boc*, *Cdon*, *Smo*, *Ptch1*, *Gli1*, *Gli2*, and *Gli3* in chick fibroblasts (i.e., DF-1) 24 hours (h) after treatment with five serial dilutions of doxycycline (dox) and induction of a stably integrated chick (yellow) or quail (green) *Gas1* overexpression vector. **(B)** Relative levels of mRNA expression for *Boc*, *Cdon*, *Smo*, *Gli2*, and *Gli3* in chick fibroblasts 48 hours after dox-induction of a stably integrated *Gas1* overexpression vector (serial dilution) and treatment with recombinant (r) SHH protein (control 0 ng/mL in white and 100 ng/mL dark grey). Expression levels were assayed by qRT-PCR and normalized to *r18s*. Significance is shown for comparisons between control (0 ng/mL of rSHH and/or dox) and treatment groups as denoted by brackets and p-values are as indicated. P-values ≥ 0.05 are considered not significant. n ≥ 2 for each group and data point, and error bars represent SEM.

**Supplemental Table 1.**
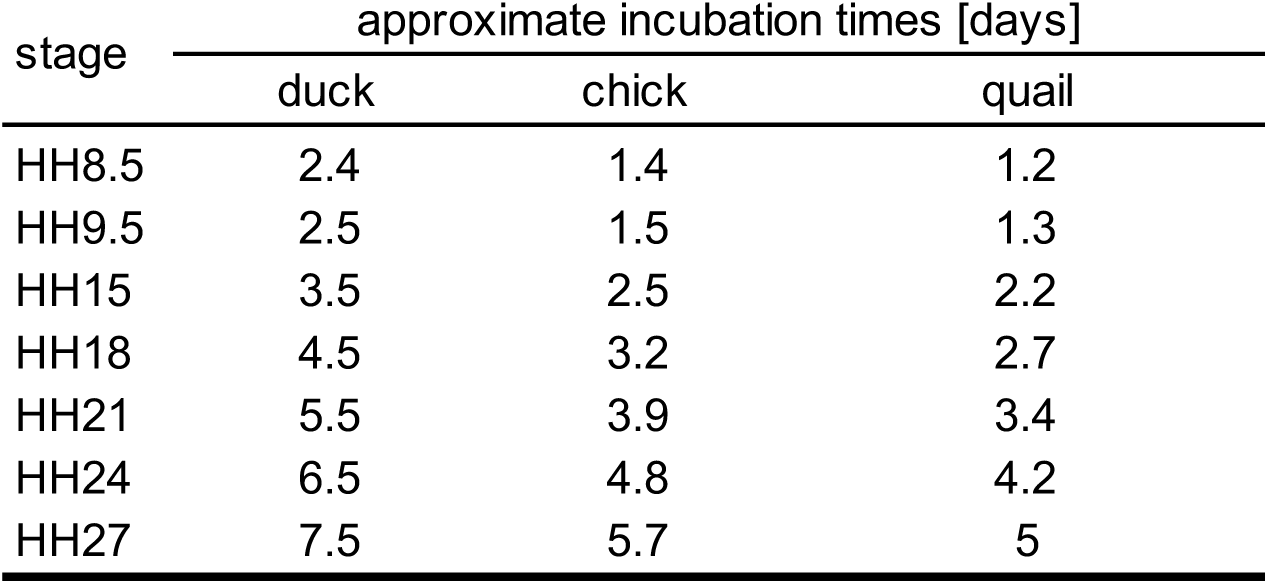
Duck, chick, and quail incubation times.

**Supplemental Table 2.** Mandibular mesenchyme population size in duck, chick, and quail embryos.

| stage | duck |  |  | chick |  |  | quail |  |  |
| --- | --- | --- | --- | --- | --- | --- | --- | --- | --- |
|  | mean | SEM | n | mean | SEM | n | mean | SEM | n |
| HH18 | 68206.8 | 10378.0 | 9 | 30296.3 | 2154.3 | 9 | 18574.7 | 3991.7 | 9 |
| HH21 | 279777.8 | 24087.8 | 12 | 119858.6 | 6436.9 | 11 | 59793.2 | 4784.6 | 9 |
| HH24 | 638656.6 | 26323.9 | 11 | 305370.4 | 15696.2 | 6 | 163454.6 | 14439.6 | 11 |
| HH27 | 2105453.7 | 74286.0 | 18 | 853886.0 | 40050.0 | 11 | 505857.1 | 42977.0 | 7 |

**Supplemental Table 3.** P-values for mandibular mesenchyme population size (significant values highlighted in grey)

| stage | mandibular mesenchyme population size |  |  |  |  |
| --- | --- | --- | --- | --- | --- |
|  | duck vs chick | duck vs quail | quail vs chick | duck vs duck<br>*Gas1* OE | quail vs quail<br>*Gas1* OE |
| HH18 | 2.52E-03 | 3.92E-04 | 2.00E-02 | 8.34E-01 | 6.14E-02 |
| HH21 | 8.18E-06 | 5.11E-07 | 4.27E-06 | 2.99E-05 | 5.70E-02 |
| HH24 | 7.89E-07 | 3.56E-12 | 4.93E-05 | 1.96E-04 | 6.14E-02 |
| HH27 | 4.31E-12 | 1.41E-11 | 6.41E-05 | - | - |

**Supplemental Table 4.** Duck/quail mandibular mesenchyme population size ratios.

| stage | HH18 | HH21 | HH24 | HH27 |
| --- | --- | --- | --- | --- |
| in vivo | 3.67 | 4.68 | 3.91 | 4.16 |
| model | 3.61 | 4.41 | 4.29 | 4.17 |

**Supplemental Table 5.**
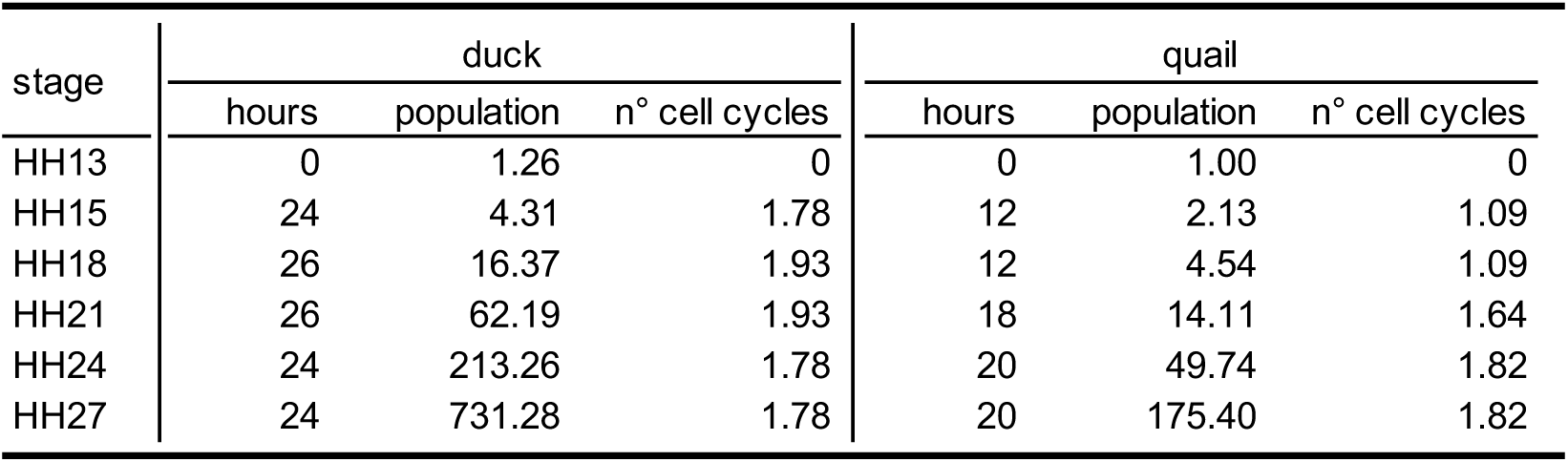
Mandibular mesenchyme population size modeling in duck and quail embryos.

| stage | duck |  |  | quail |  |  |
| --- | --- | --- | --- | --- | --- | --- |
|  | hours | population | n° cell cycles | hours | population | n° cell cycles |
| HH13 | 0 | 1.26 | 0 | 0 | 1.00 | 0 |
| HH15 | 24 | 4.31 | 1.78 | 12 | 2.13 | 1.09 |
| HH18 | 26 | 16.37 | 1.93 | 12 | 4.54 | 1.09 |
| HH21 | 26 | 62.19 | 1.93 | 18 | 14.11 | 1.64 |
| HH24 | 24 | 213.26 | 1.78 | 20 | 49.74 | 1.82 |
| HH27 | 24 | 731.28 | 1.78 | 20 | 175.40 | 1.82 |

**Supplemental Table 6.**
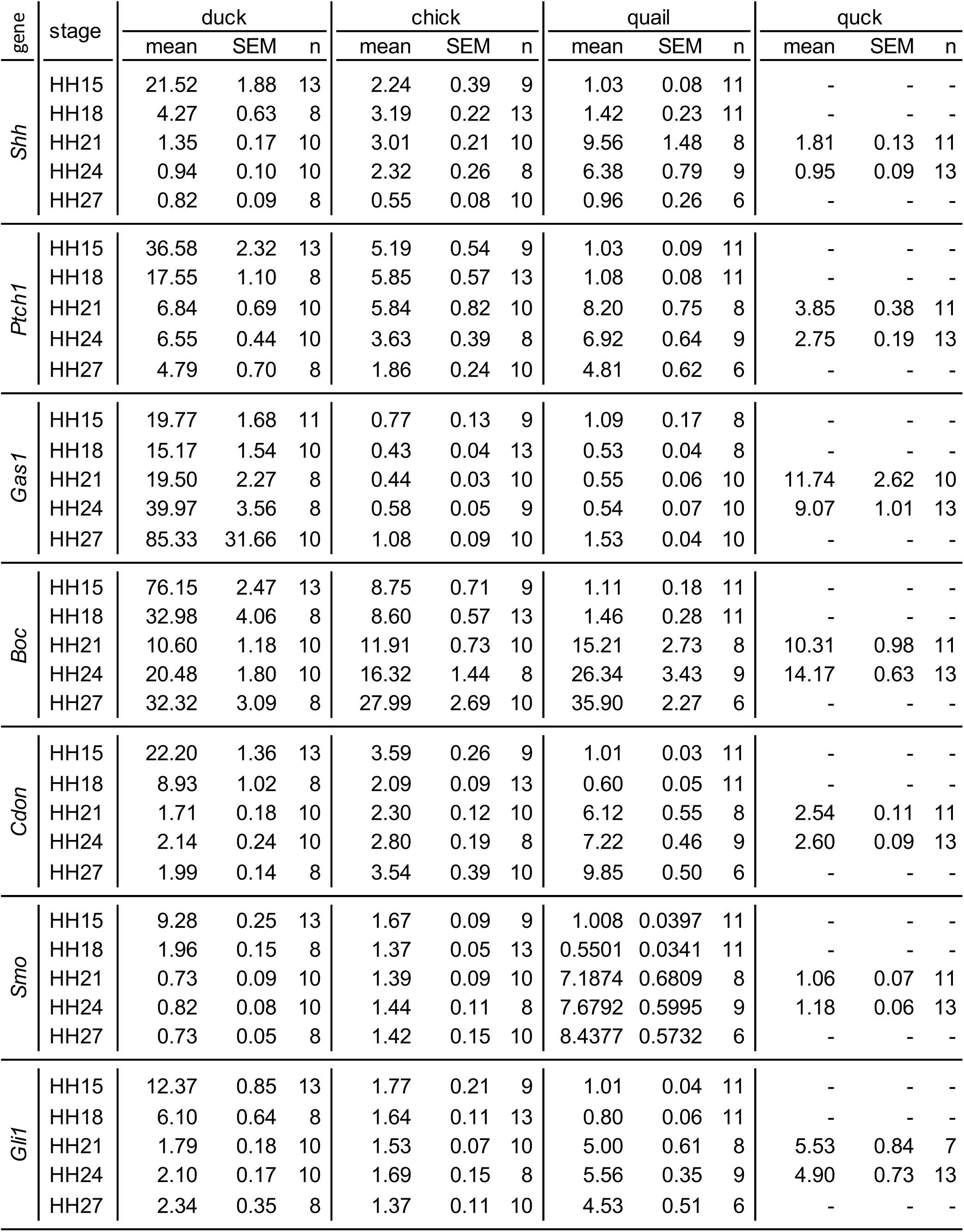
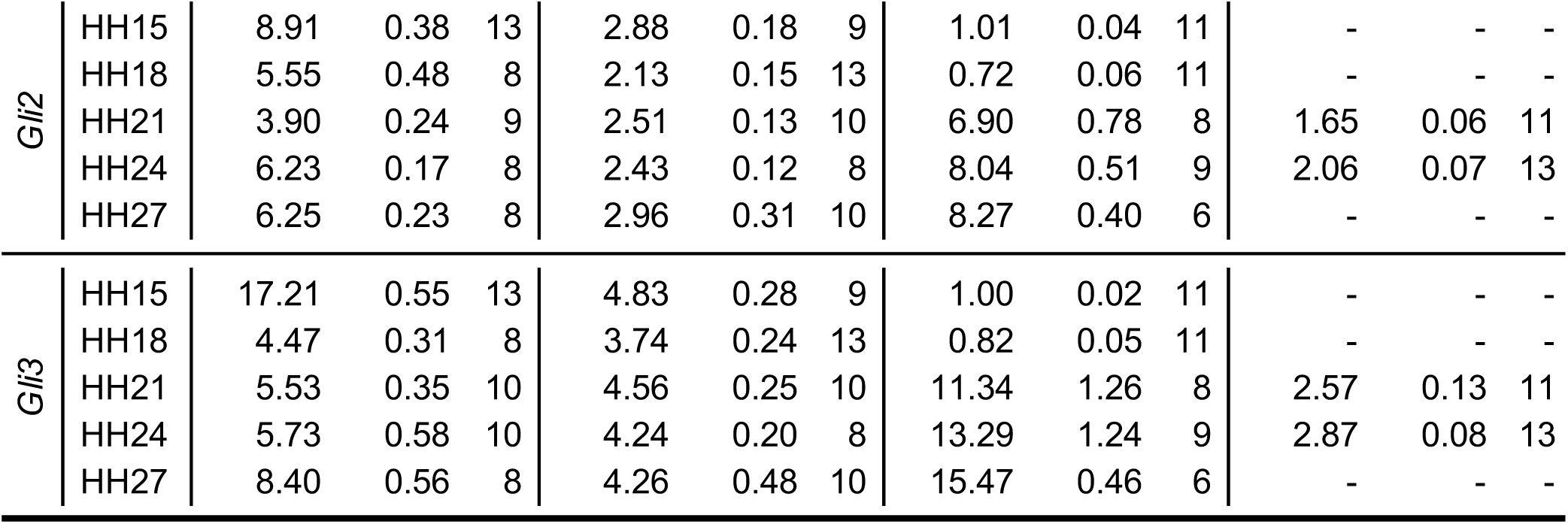
SHH pathway members mRNA levels in duck, chick, quail, and quck mandibular primordia.

**Supplemental Table 7.** Shh pathway gene expression ratios in duck, chick, quail, and quck mandibular primordia.

|  | stage | duck |  |  | chick |  |  | quail |  |  | quck |  |  |
| --- | --- | --- | --- | --- | --- | --- | --- | --- | --- | --- | --- | --- | --- |
|  |  | mean | SEM | n | mean | SEM | n | mean | SEM | n | mean | SEM | n |
| *Shh/Ptch1* | HH15 | 0.62 | 0.07 | 13 | 0.41 | 0.04 | 9 | 1.06 | 0.12 | 11 | - | - | - |
|  | HH18 | 0.25 | 0.04 | 8 | 0.59 | 0.06 | 13 | 1.37 | 0.24 | 11 | - | - | - |
|  | HH21 | 0.21 | 0.03 | 10 | 0.61 | 0.09 | 10 | 1.07 | 0.20 | 7 | 0.50 | 0.05 | 11 |
|  | HH24 | 0.14 | 0.01 | 10 | 0.65 | 0.05 | 8 | 1.03 | 0.19 | 9 | 0.35 | 0.03 | 13 |
|  | HH27 | 0.18 | 0.02 | 8 | 0.31 | 0.04 | 10 | 0.21 | 0.05 | 6 | - | - | - |
| *Shh/Gli1* | HH15 | 1.89 | 0.25 | 13 | 1.22 | 0.12 | 9 | 1.03 | 0.09 | 11 | - | - | - |
|  | HH18 | 0.71 | 0.10 | 8 | 1.99 | 0.14 | 13 | 1.99 | 0.38 | 11 | - | - | - |
|  | HH21 | 0.78 | 0.09 | 10 | 1.98 | 0.11 | 10 | 1.73 | 0.37 | 7 | 0.26 | 0.05 | 10 |
|  | HH24 | 0.45 | 0.04 | 10 | 1.39 | 0.13 | 8 | 1.23 | 0.20 | 9 | 0.25 | 0.05 | 13 |
|  | HH27 | 0.39 | 0.06 | 8 | 0.40 | 0.05 | 10 | 0.22 | 0.06 | 6 | - | - | - |
| *Shh/Gli2* | HH15 | 2.46 | 0.23 | 13 | 0.78 | 0.13 | 9 | 1.02 | 0.07 | 11 | - | - | - |
|  | HH18 | 0.77 | 0.09 | 8 | 1.53 | 0.10 | 13 | 1.92 | 0.26 | 11 | - | - | - |
|  | HH21 | 0.36 | 0.06 | 9 | 1.21 | 0.09 | 10 | 1.59 | 0.41 | 8 | 1.12 | 0.12 | 11 |
|  | HH24 | 0.14 | 0.01583 | 8 | 2.43 | 0.12 | 8 | 0.81 | 0.11 | 9 | 0.47 | 0.05 | 13 |
|  | HH27 | 0.13 | 0.02 | 8 | 2.96 | 0.31 | 10 | 0.11 | 0.03 | 6 | - | - | - |
| *Shh/Gli3* | HH15 | 1.26 | 0.12 | 13 | 0.47 | 0.08 | 9 | 1.03 | 0.08 | 11 | - | - | - |
|  | HH18 | 0.94 | 0.11 | 8 | 0.87 | 0.06 | 13 | 1.86 | 0.37 | 11 | - | - | - |
|  | HH21 | 0.25 | 0.03 | 10 | 0.67 | 0.06 | 10 | 1.08 | 0.36 | 8 | 0.74 | 0.09 | 11 |
|  | HH24 | 0.18 | 0.03 | 10 | 0.56 | 0.07 | 8 | 0.52 | 0.08 | 9 | 0.34 | 0.04 | 13 |
|  | HH27 | 0.10 | 0.01 | 8 | 0.13 | 0.02 | 10 | 0.06 | 0.02 | 6 | - | - | - |

**Supplemental Table 8.** P-values for in vivo samples (significant values highlighted in grey)

| stage | *Shh* mRNA level |  |  |
| --- | --- | --- | --- |
|  | duck vs chick | duck vs quail | quail vs chick |
| HH15 | < 1.00E-06 | < 1.00E-06 | 7.55E-03 |
| HH18 | 8.92E-02 | 3.61E-04 | 8.80E-05 |
| HH21 | 3.40E-05 | 4.00E-05 | 6.50E-04 |
| HH24 | 1.57E-04 | 6.00E-06 | 9.44E-04 |
| HH27 | 8.92E-02 | 5.77E-01 | 9.22E-02 |

| stage | *Boc* mRNA level |  |  |
| --- | --- | --- | --- |
|  | duck vs chick | duck vs quail | quail vs chick |
| HH15 | < 1.00E-06 | < 1.00E-06 | < 1.00E-06 |
| HH18 | < 1.00E-06 | < 1.00E-06 | < 1.00E-06 |
| HH21 | 5.16E-01 | 3.06E-01 | 2.15E-01 |
| HH24 | 2.76E-01 | 3.06E-01 | 6.21E-02 |
| HH27 | 5.16E-01 | 3.99E-01 | 1.22E-01 |

| stage | *Cdon* mRNA level |  |  |
| --- | --- | --- | --- |
|  | duck vs chick | duck vs quail | quail vs chick |
| HH15 | < 1.00E-06 | < 1.00E-06 | < 1.00E-06 |
| HH18 | < 1.00E-06 | < 1.00E-06 | < 1.00E-06 |
| HH21 | 2.88E-02 | < 1.00E-06 | 1.00E-06 |
| HH24 | 5.76E-02 | < 1.00E-06 | < 1.00E-06 |
| HH27 | 1.04E-02 | < 1.00E-06 | < 1.00E-06 |

| stage | *Gas1* mRNA level |  |  |
| --- | --- | --- | --- |
|  | duck vs chick | duck vs quail | quail vs chick |
| HH15 | < 1.00E-06 | < 1.00E-06 | 4.06E-01 |
| HH18 | < 1.00E-06 | < 1.00E-06 | 4.06E-01 |
| HH21 | < 1.00E-06 | < 1.00E-06 | 4.06E-01 |
| HH24 | < 1.00E-06 | < 1.00E-06 | 6.43E-01 |
| HH27 | 1.59E-02 | 1.64E-02 | 1.19E-03 |

| stage | *Ptch1* mRNA level |  |  |
| --- | --- | --- | --- |
|  | duck vs chick | duck vs quail | quail vs chick |
| HH15 | < 1.00E-06 | < 1.00E-06 | < 1.00E-06 |
| HH18 | < 1.00E-06 | < 1.00E-06 | < 1.00E-06 |
| HH21 | 3.63E-01 | 4.90E-01 | 5.49E-02 |
| HH24 | 5.42E-04 | 8.68E-01 | 1.46E-03 |
| HH27 | 1.01E-03 | 9.79E-01 | 3.75E-04 |

| stage | *Smo* mRNA level |  |  |
| --- | --- | --- | --- |
|  | duck vs chick | duck vs quail | quail vs chick |
| HH15 | < 1.00E-06 | < 1.00E-06 | < 1.00E-06 |
| HH18 | 7.60E-04 | < 1.00E-06 | < 1.00E-06 |
| HH21 | 1.76E-04 | < 1.00E-06 | < 1.00E-06 |
| HH24 | 6.69E-04 | < 1.00E-06 | < 1.00E-06 |
| HH27 | 1.35E-03 | < 1.00E-06 | < 1.00E-06 |

| stage | *Gli1* mRNA level |  |  |
| --- | --- | --- | --- |
|  | duck vs chick | duck vs quail | quail vs chick |
| HH15 | < 1.00E-06 | < 1.00E-06 | 8.40E-04 |
| HH18 | < 1.00E-06 | < 1.00E-06 | 7.00E-06 |
| HH21 | 1.95E-01 | 9.1E-05 | 2.10E-05 |
| HH24 | 1.93E-01 | < 1.00E-06 | < 1.00E-06 |
| HH27 | 3.09E-02 | 0.00336 | 7.00E-06 |

| stage | *Gli2* mRNA level |  |  |
| --- | --- | --- | --- |
|  | duck vs chick | duck vs quail | quail vs chick |
| HH15 | < 1.00E-06 | < 1.00E-06 | < 1.00E-06 |
| HH18 | < 1.00E-06 | < 1.00E-06 | < 1.00E-06 |
| HH21 | 7.9E-05 | 3.00E-03 | 1.20E-05 |
| HH24 | < 1.00E-06 | 5.63E-03 | < 1.00E-06 |
| HH27 | < 1.00E-06 | 1.76E-03 | < 1.00E-06 |

| stage | *Gli3* mRNA level |  |  |
| --- | --- | --- | --- |
|  | duck vs chick | duck vs quail | quail vs chick |
| HH15 | < 1.00E-06 | < 1.00E-06 | < 1.00E-06 |
| HH18 | 1.04E-01 | < 1.00E-06 | < 1.00E-06 |
| HH21 | 1.04E-01 | 1.60E-04 | 2.40E-05 |
| HH24 | 1.04E-01 | 5.00E-05 | 1.20E-05 |
| HH27 | 1.35E-04 | 2.00E-06 | < 1.00E-06 |

| stage | *Shh* to *Ptch1* ratios |  |  |
| --- | --- | --- | --- |
|  | duck vs chick | duck vs quail | quail vs chick |
| HH15 | 4.73E-02 | 5.91E-03 | 6.76E-04 |
| HH18 | 1.59E-03 | 3.49E-03 | 9.72E-03 |
| HH21 | 2.57E-03 | 6.60E-04 | 1.10E-01 |
| HH24 | < 1.00E-06 | 5.53E-04 | 1.64E-01 |
| HH27 | 4.73E-02 | 6.16E-01 | 1.68E-01 |

| stage | *Shh* to *Gli1* ratios |  |  |
| --- | --- | --- | --- |
|  | duck vs chick | duck vs quail | quail vs chick |
| HH15 | 1.01E-01 | 2.79E-02 | 6.02E-01 |
| HH18 | 6.00E-06 | 2.93E-02 | 9.94E-01 |
| HH21 | < 1.00E-06 | 2.93E-02 | 8.53E-01 |
| HH24 | 3.00E-06 | 4.83E-03 | 8.53E-01 |
| HH27 | 9.51E-01 | 5.43E-02 | 1.54E-01 |

**Supplemental Table 9.** SHH pathway members mRNA levels in duck, chick, and quail mandibular primordia explant cultures.

| gene | treatment | duck |  |  | chick |  |  | quail |  |  |
| --- | --- | --- | --- | --- | --- | --- | --- | --- | --- | --- |
|  |  | mean | SEM | n | mean | SEM | n | mean | SEM | n |
| *Shh* | ctrl | 1.14 | 0.14 | 17 | 1.45 | 0.25 | 12 | 1.06 | 0.10 | 14 |
|  | cycl | 1.78 | 0.25 | 8 | 2.11 | 0.35 | 8 | 1.55 | 0.23 | 8 |
|  | 0.1 | 0.76 | 0.27 | 4 | 1.60 | 0.30 | 7 | 0.71 | 0.07 | 7 |
|  | 1 | 0.54 | 0.06 | 6 | 1.24 | 0.33 | 7 | 1.14 | 0.28 | 7 |
|  | 10 | 0.47 | 0.04 | 6 | 0.54 | 0.10 | 7 | 0.84 | 0.11 | 6 |
|  | 100 | 0.53 | 0.09 | 6 | 0.69 | 0.25 | 7 | 0.66 | 0.07 | 7 |
|  | 1000 | 0.57 | 0.16 | 13 | 0.53 | 0.19 | 7 | 0.47 | 0.13 | 8 |
| *Ptch1* | ctrl | 1.04 | 0.08 | 17 | 1.04 | 0.08 | 12 | 1.04 | 0.08 | 14 |
|  | cycl | 0.23 | 0.02 | 8 | 0.12 | 0.01 | 8 | 0.15 | 0.01 | 8 |
|  | 0.1 | 0.83 | 0.18 | 4 | 0.93 | 0.09 | 7 | 0.77 | 0.09 | 7 |
|  | 1 | 0.83 | 0.10 | 6 | 0.93 | 0.16 | 7 | 0.84 | 0.13 | 7 |
|  | 10 | 0.87 | 0.13 | 6 | 1.16 | 0.17 | 7 | 0.86 | 0.04 | 6 |
|  | 100 | 1.39 | 0.14 | 6 | 1.33 | 0.16 | 7 | 1.17 | 0.13 | 7 |
|  | 1000 | 2.46 | 0.25 | 13 | 3.44 | 0.44 | 7 | 1.47 | 0.12 | 8 |
| *Gas1* | ctrl | 1.08 | 0.11 | 17 | 1.06 | 0.11 | 12 | 1.07 | 0.11 | 14 |
|  | cycl | 1.93 | 0.21 | 8 | 2.25 | 0.28 | 8 | 1.14 | 0.10 | 8 |
|  | 0.1 | 1.28 | 0.05 | 4 | 0.87 | 0.10 | 7 | 0.68 | 0.08 | 7 |
|  | 1 | 1.18 | 0.14 | 6 | 0.76 | 0.11 | 7 | 0.68 | 0.05 | 7 |
|  | 10 | 1.09 | 0.24 | 6 | 0.90 | 0.06 | 7 | 0.55 | 0.07 | 6 |
|  | 100 | 0.95 | 0.11 | 6 | 0.52 | 0.04 | 7 | 0.55 | 0.04 | 7 |
|  | 1000 | 0.95 | 0.05 | 13 | 0.46 | 0.05 | 7 | 0.24 | 0.02 | 8 |
| *Boc* | ctrl | 1.02 | 0.05 | 17 | 1.01 | 0.05 | 12 | 1.04 | 0.08 | 14 |
|  | cycl | 1.06 | 0.08 | 8 | 1.00 | 0.11 | 8 | 0.87 | 0.08 | 8 |
|  | 0.1 | 1.32 | 0.10 | 4 | 0.93 | 0.08 | 7 | 0.97 | 0.06 | 7 |
|  | 1 | 1.12 | 0.05 | 6 | 1.05 | 0.07 | 7 | 0.97 | 0.08 | 7 |
|  | 10 | 1.11 | 0.10 | 6 | 1.13 | 0.07 | 7 | 0.71 | 0.07 | 6 |
|  | 100 | 0.91 | 0.14 | 6 | 0.73 | 0.04 | 7 | 0.61 | 0.06 | 7 |
|  | 1000 | 1.00 | 0.05 | 13 | 0.51 | 0.03 | 7 | 0.54 | 0.09 | 8 |
| *Cdon* | ctrl | 1.02 | 0.06 | 17 | 1.02 | 0.06 | 12 | 1.02 | 0.05 | 14 |
|  | cycl | 1.48 | 0.10 | 8 | 1.15 | 0.11 | 8 | 1.13 | 0.08 | 8 |
|  | 0.1 | 1.16 | 0.08 | 4 | 0.99 | 0.08 | 7 | 0.95 | 0.07 | 7 |
|  | 1 | 1.08 | 0.07 | 6 | 1.07 | 0.06 | 7 | 0.92 | 0.04 | 7 |
|  | 10 | 1.17 | 0.10 | 6 | 1.18 | 0.06 | 7 | 0.88 | 0.05 | 6 |
|  | 100 | 1.13 | 0.12 | 6 | 0.92 | 0.06 | 7 | 0.75 | 0.08 | 7 |
|  | 1000 | 0.94 | 0.06 | 13 | 0.65 | 0.04 | 7 | 0.48 | 0.07 | 8 |
| *Smo* | ctrl | 1.01 | 0.04 | 17 | 1.01 | 0.03 | 12 | 1.04 | 0.08 | 14 |
|  | cycl | 1.01 | 0.06 | 8 | 0.90 | 0.05 | 8 | 0.82 | 0.08 | 8 |
|  | 0.1 | 1.16 | 0.11 | 4 | 0.96 | 0.05 | 7 | 0.71 | 0.07 | 7 |
|  | 1 | 1.12 | 0.08 | 6 | 0.92 | 0.04 | 7 | 0.84 | 0.07 | 7 |
|  | 10 | 1.04 | 0.06 | 6 | 0.98 | 0.06 | 7 | 0.80 | 0.06 | 6 |
|  | 100 | 1.05 | 0.09 | 6 | 0.93 | 0.06 | 7 | 0.83 | 0.06 | 7 |
|  | 1000 | 1.16 | 0.05 | 13 | 0.84 | 0.07 | 7 | 0.77 | 0.13 | 8 |
| *Gli1* | ctrl | 1.03 | 0.06 | 17 | 1.02 | 0.06 | 12 | 1.04 | 0.08 | 14 |
|  | cycl | 0.21 | 0.03 | 8 | 0.12 | 0.03 | 8 | 0.17 | 0.02 | 8 |
|  | 0.1 | 1.42 | 0.33 | 4 | 0.96 | 0.05 | 7 | 0.78 | 0.10 | 7 |
|  | 1 | 1.33 | 0.13 | 6 | 1.02 | 0.05 | 7 | 0.76 | 0.05 | 7 |
|  | 10 | 1.26 | 0.13 | 6 | 1.04 | 0.05 | 7 | 0.89 | 0.05 | 6 |
|  | 100 | 1.57 | 0.19 | 6 | 1.18 | 0.08 | 7 | 0.83 | 0.05 | 7 |
|  | 1000 | 1.84 | 0.16 | 13 | 1.62 | 0.09 | 7 | 1.07 | 0.11 | 8 |
| *Gli2* | ctrl | 1.01 | 0.04 | 17 | 1.01 | 0.03 | 12 | 1.02 | 0.05 | 14 |
|  | cycl | 0.81 | 0.06 | 8 | 0.99 | 0.05 | 8 | 0.69 | 0.03 | 8 |
|  | 0.1 | 1.24 | 0.06 | 4 | 0.88 | 0.03 | 7 | 0.98 | 0.07 | 7 |
|  | 1 | 1.12 | 0.06 | 6 | 0.92 | 0.06 | 7 | 0.90 | 0.05 | 7 |
|  | 10 | 1.06 | 0.08 | 6 | 0.95 | 0.04 | 7 | 0.95 | 0.02 | 6 |
|  | 100 | 1.10 | 0.07 | 6 | 0.82 | 0.04 | 7 | 0.84 | 0.03 | 7 |
|  | 1000 | 1.18 | 0.07 | 13 | 0.76 | 0.05 | 7 | 0.70 | 0.04 | 8 |
| *Gli3* | ctrl | 1.03 | 0.06 | 17 | 1.01 | 0.03 | 12 | 1.02 | 0.05 | 14 |
|  | cycl | 1.27 | 0.08 | 8 | 0.92 | 0.04 | 8 | 0.79 | 0.05 | 8 |
|  | 0.1 | 1.30 | 0.22 | 4 | 1.00 | 0.05 | 7 | 0.94 | 0.06 | 7 |
|  | 1 | 1.08 | 0.07 | 6 | 0.92 | 0.04 | 7 | 0.89 | 0.05 | 7 |
|  | 10 | 1.09 | 0.03 | 6 | 0.98 | 0.04 | 7 | 0.81 | 0.04 | 6 |
|  | 100 | 0.97 | 0.05 | 6 | 0.76 | 0.04 | 7 | 0.73 | 0.02 | 7 |
|  | 1000 | 0.89 | 0.05 | 13 | 0.58 | 0.02 | 7 | 0.53 | 0.06 | 8 |

**Supplemental Table 10.**
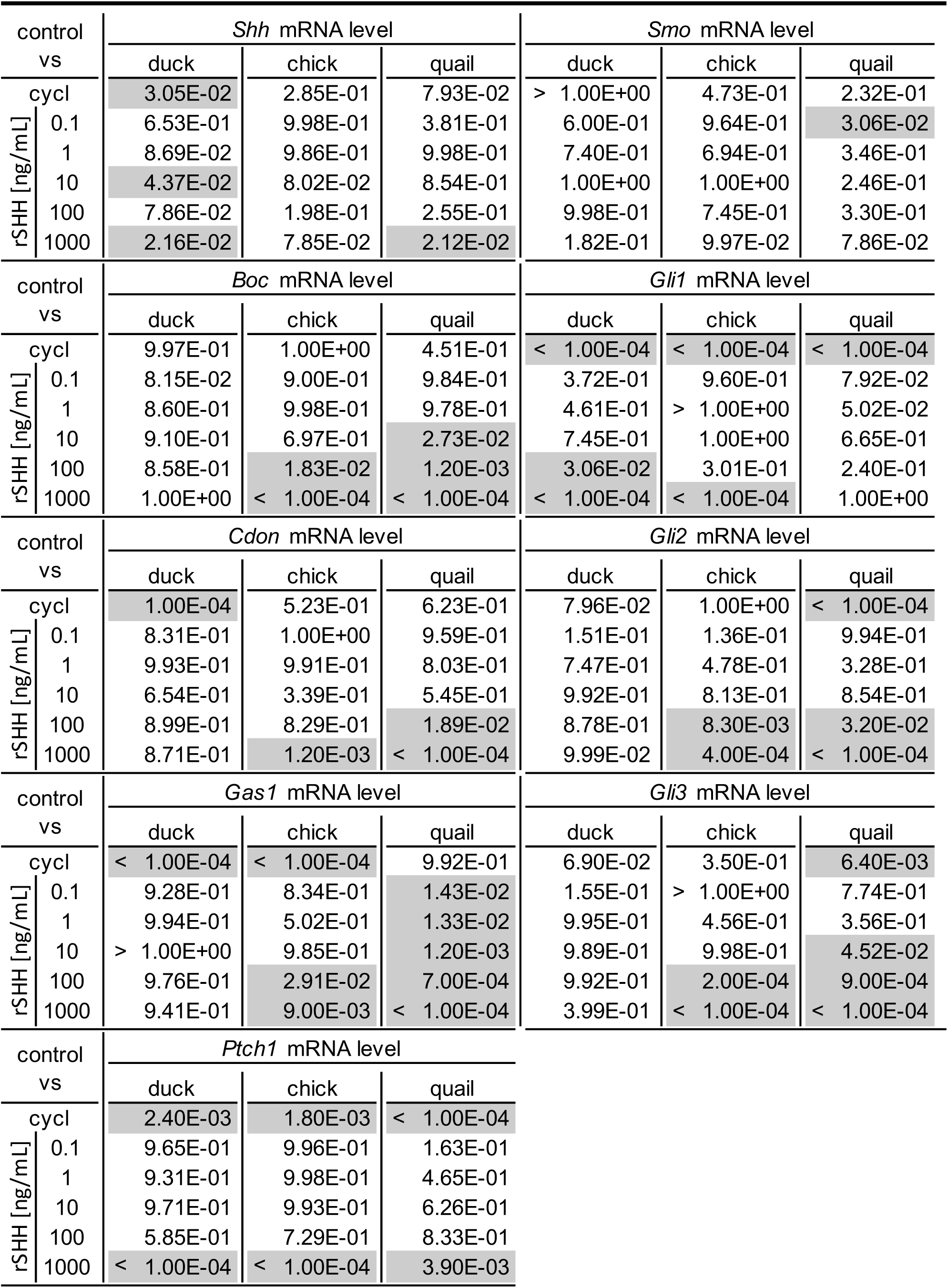
P-values for mandibular primordia explant cultures (significant values in grey)

**Supplemental Table 11.** P-values for in vivo samples - quck data (significant values highlighted in grey)

| quck stage | *Shh* mRNA level |  |  |
| --- | --- | --- | --- |
|  | duck vs quail | duck vs quck | quail vs quck |
| HH21 | < 1.00E-04 | 5.54E-01 | < 1.00E-04 |
| HH24 | 9.99E-01 | 1.00E+00 | 1.00E+00 |
| quck stage | *Boc* mRNA level |  |  |
|  | duck vs quail | duck vs quck | quail vs quck |
| HH21 | < 1.00E-04 | 9.92E-01 | < 1.00E-04 |
| HH24 | < 1.00E-04 | 2.24E-02 | < 1.00E-04 |
| quck stage | *Cdon* mRNA level |  |  |
|  | duck vs quail | duck vs quck | quail vs quck |
| HH21 | < 1.00E-04 | 5.37E-02 | < 1.00E-04 |
| HH24 | < 1.00E-04 | 3.65E-01 | < 1.00E-04 |
| quck stage | *Gas1* mRNA level |  |  |
|  | duck vs quail | duck vs quck | quail vs quck |
| HH21 | < 1.00E-04 | 1.80E-02 | 2.00E-04 |
| HH24 | < 1.00E-04 | < 1.00E-04 | 8.60E-03 |
| quck stage | *Smo* mRNA level |  |  |
|  | duck vs quail | duck vs quck | quail vs quck |
| HH21 | < 1.00E-04 | 6.48E-01 | < 1.00E-04 |
| HH24 | < 1.00E-04 | 5.65E-01 | < 1.00E-04 |
| quck stage | *Gli1* mRNA level |  |  |
|  | duck vs quail | duck vs quck | quail vs quck |
| HH21 | < 1.00E-04 | < 1.00E-04 | 9.99E-01 |
| HH24 | 1.81E-02 | 6.00E-04 | 8.91E-01 |
| quck stage | *Shh* to *Ptch1* ratios |  |  |
|  | duck vs quail | duck vs quck | quail vs quck |
| HH21 | < 1.00E-04 | 2.37E-02 | < 1.00E-04 |
| HH24 | 8.50E-01 | 1.19E-01 | 4.90E-01 |
| quck stage | *Shh* to *Gli2* ratios |  |  |
|  | duck vs quail | duck vs quck | quail vs quck |
| HH21 | 9.00E-04 | < 1.00E-04 | 2.06E-02 |
| HH24 | 9.82E-01 | 1.22E-02 | 1.45E-02 |
| quck stage | *Shh* to *Gli3* ratios |  |  |
|  | duck vs quail | duck vs quck | quail vs quck |
| HH21 | 7.00E-03 | < 1.00E-04 | 2.36E-02 |
| HH24 | 4.47E-01 | 1.08E-01 | 1.03E-02 |

**Supplemental Table 12.** SHH pathway members mRNA levels in DF1 cells after 24 hours of recombinant SHH (rSHH) treatment.

| rSHH | *Gas1* |  |  | *Ptch1* |  |  | *Gli1* |  |  |
| --- | --- | --- | --- | --- | --- | --- | --- | --- | --- |
|  | mean | SEM | n | mean | SEM | n | mean | SEM | n |
| 0 | 1.00 | 0.03 | 8 | 1.00 | 0.02 | 8 | 1.06 | 0.12 | 8 |
| 0.1 | 0.96 | 0.07 | 8 | 0.90 | 0.06 | 8 | 1.03 | 0.12 | 8 |
| 1 | 1.00 | 0.06 | 8 | 1.01 | 0.08 | 8 | 1.25 | 0.23 | 8 |
| 10 | 0.91 | 0.05 | 8 | 1.08 | 0.08 | 8 | 1.53 | 0.23 | 8 |
| 100 | 0.80 | 0.05 | 8 | 1.07 | 0.05 | 8 | 1.38 | 0.28 | 8 |
| 1000 | 0.88 | 0.08 | 8 | 1.32 | 0.11 | 8 | 1.59 | 0.28 | 8 |

**Supplemental Table 13.**
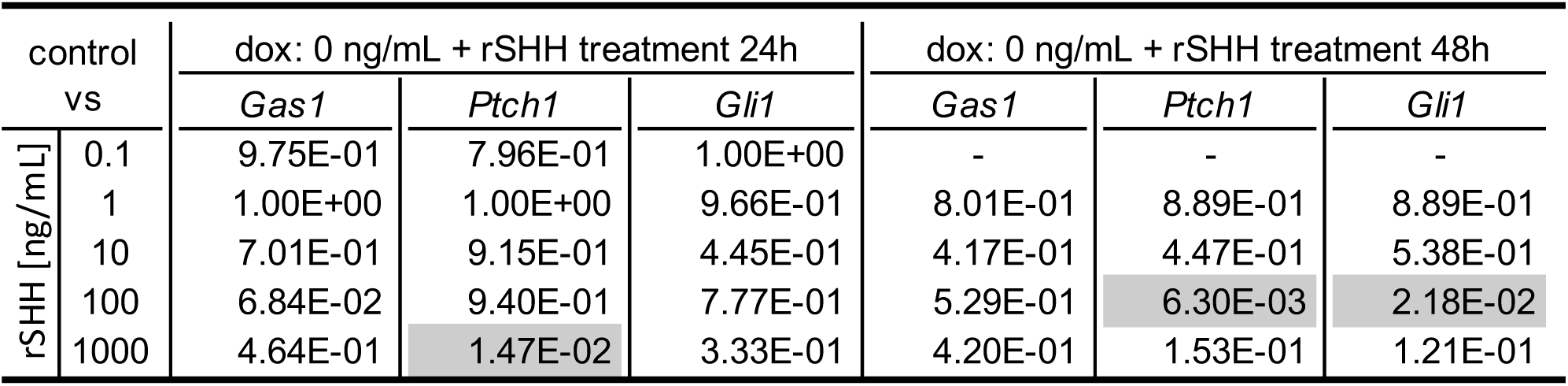
P-values for DF1 cell cultures with stably integrated dox-inducible *Gas1* construct after 24 and 48 hours of recombinant SHH (rSHH) protein treatment (significant values highlighted in gray)

**Supplemental Table 14.**
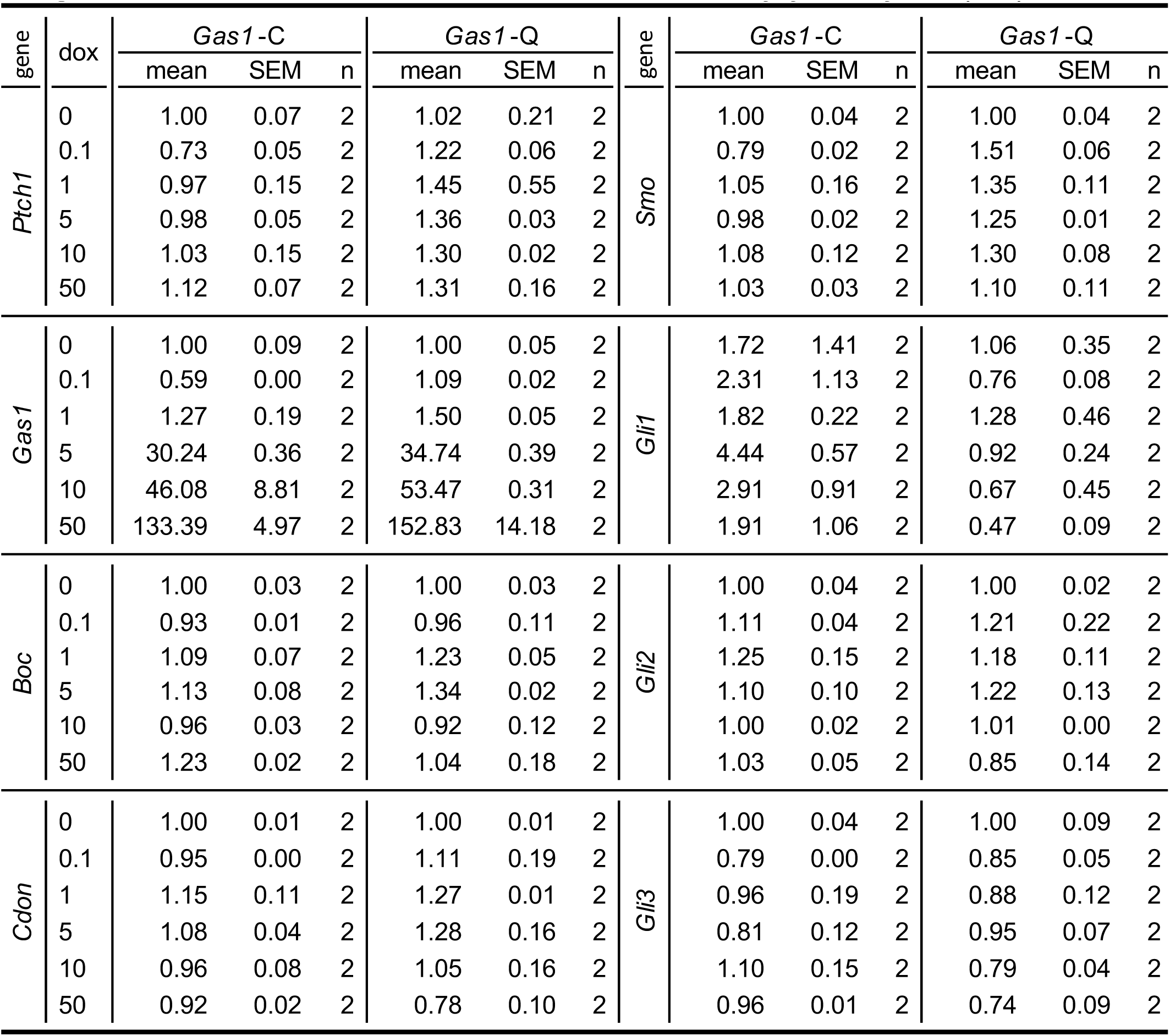
SHH pathway members mRNA levels in DF1 cell cultures with stably integrated dox-inducible *Gas1* construct after 24 hours of doxycycline hyclate (dox) treatment.

**Supplemental Table 15.** Number of cells counted in 6-well plates after 72 hours of doxycycline hyclate (dox) treatment in DF1 cell lines with stably integrated dox-inducible *Gas1*.

| dox | *Gas1*-C |  |  | *Gas1*-Q |  |  |
| --- | --- | --- | --- | --- | --- | --- |
|  | mean | SEM | n | mean | SEM | n |
| 0 | 674340.30 | 79731.16 | 4 | 404374.98 | 19182.23 | 4 |
| 0.1 | 687777.78 | 86256.14 | 4 | 447326.40 | 63572.87 | 4 |
| 1 | 690486.13 | 86948.45 | 4 | 355312.50 | 13432.59 | 4 |
| 5 | 756979.18 | 58289.62 | 4 | 354409.73 | 14914.06 | 4 |
| 10 | 593690.48 | 24880.28 | 4 | 308730.18 | 26763.42 | 4 |
| 50 | 611493.05 | 42838.07 | 4 | 322118.08 | 53244.62 | 4 |

**Supplemental Table 16.** P-values for DF1 cell cultures with stably integrated dox-inducible *Gas1* construct after 24 hours of doxycycline hyclate (dox) treatment (significant values highlighted in gray)

| gene | dox<br>0 vs | *Gas1*-C | *Gas1*-Q | gene | *Gas1*-C | *Gas1*-Q |
| --- | --- | --- | --- | --- | --- | --- |
| *Ptch1* | 0.1 | 0.3069 | 0.9623 | *Smo* | 0.3409 | 0.0125 |
|  | 1 | 0.9985 | 0.6521 |  | 0.9858 | 0.0608 |
|  | 5 | 0.9998 | 0.7991 |  | 0.9997 | 0.1932 |
|  | 10 | 0.9997 | 0.8881 |  | 0.9475 | 0.1054 |
|  | 50 | 0.8656 | 0.8783 |  | 0.9997 | 0.8438 |
| *Gas1* | 0.1 | > 0.9999 | > 0.9999 | *Gli1* | 0.968 | 0.9309 |
|  | 1 | > 0.9999 | > 0.9999 |  | 0.9997 | 0.9797 |
|  | 5 | 0.0088 | 0.0218 |  | 0.1994 | 0.9968 |
|  | 10 | 0.0009 | 0.0025 |  | 0.748 | 0.8538 |
|  | 50 | < 0.0001 | < 0.0001 |  | 0.9986 | 0.5902 |
| *Boc* | 0.1 | 0.7323 | 0.9986 | *Gli2* | 0.7634 | 0.6894 |
|  | 1 | 0.6035 | 0.4374 |  | 0.1974 | 0.7809 |
|  | 5 | 0.2995 | 0.1702 |  | 0.8542 | 0.6577 |
|  | 10 | 0.9523 | 0.9694 |  | > 0.9999 | > 0.9999 |
|  | 50 | 0.0515 | 0.9974 |  | 0.9971 | 0.8717 |
| *Cdon* | 0.1 | 0.9357 | 0.9483 | *Gli3* | 0.5971 | 0.5847 |
|  | 1 | 0.3499 | 0.4978 |  | 0.9981 | 0.7224 |
|  | 5 | 0.7706 | 0.4512 |  | 0.6611 | 0.9744 |
|  | 10 | 0.9844 | 0.9985 |  | 0.954 | 0.3125 |
|  | 50 | 0.7565 | 0.6495 |  | 0.9984 | 0.1895 |

**Supplemental Table 17.**
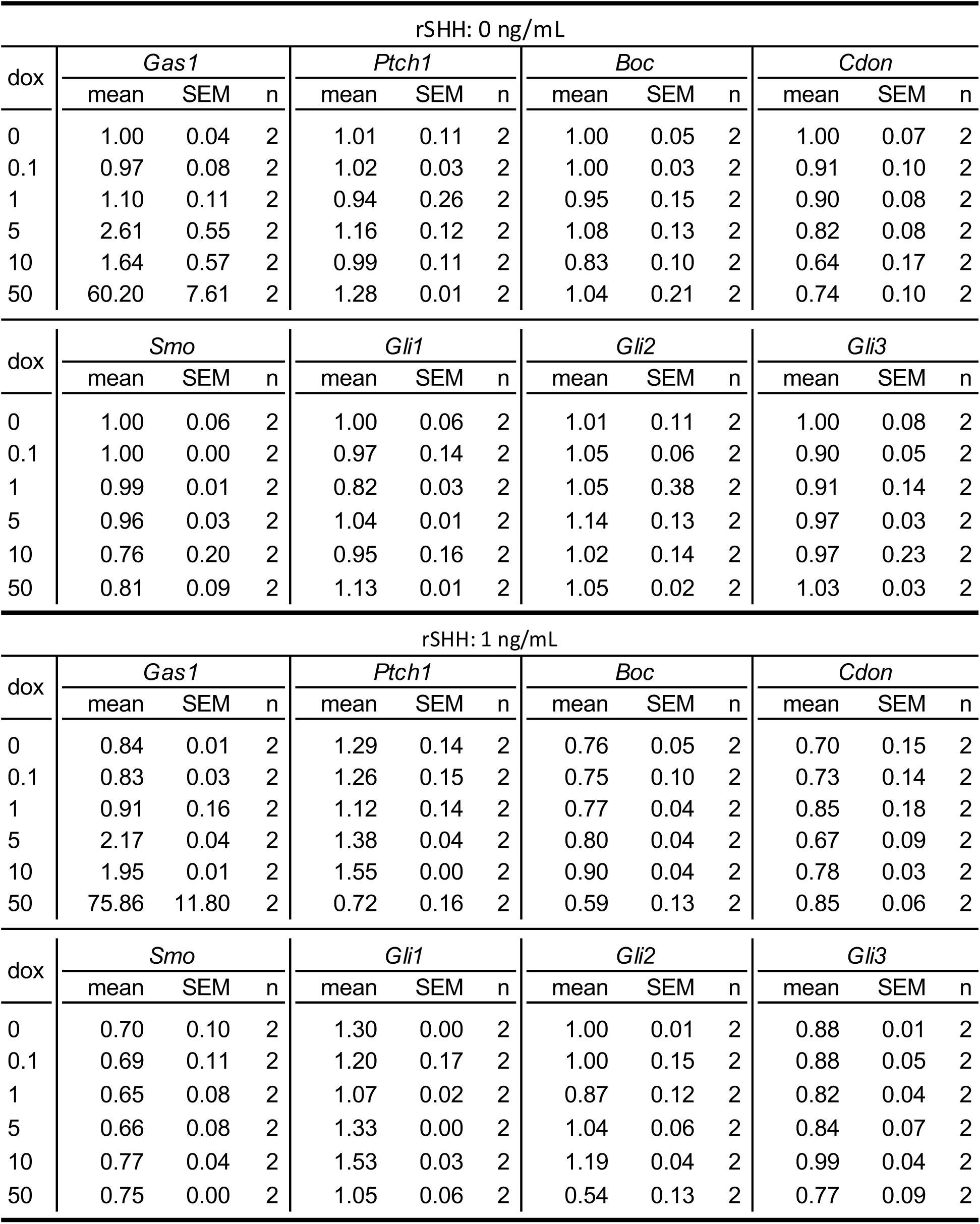
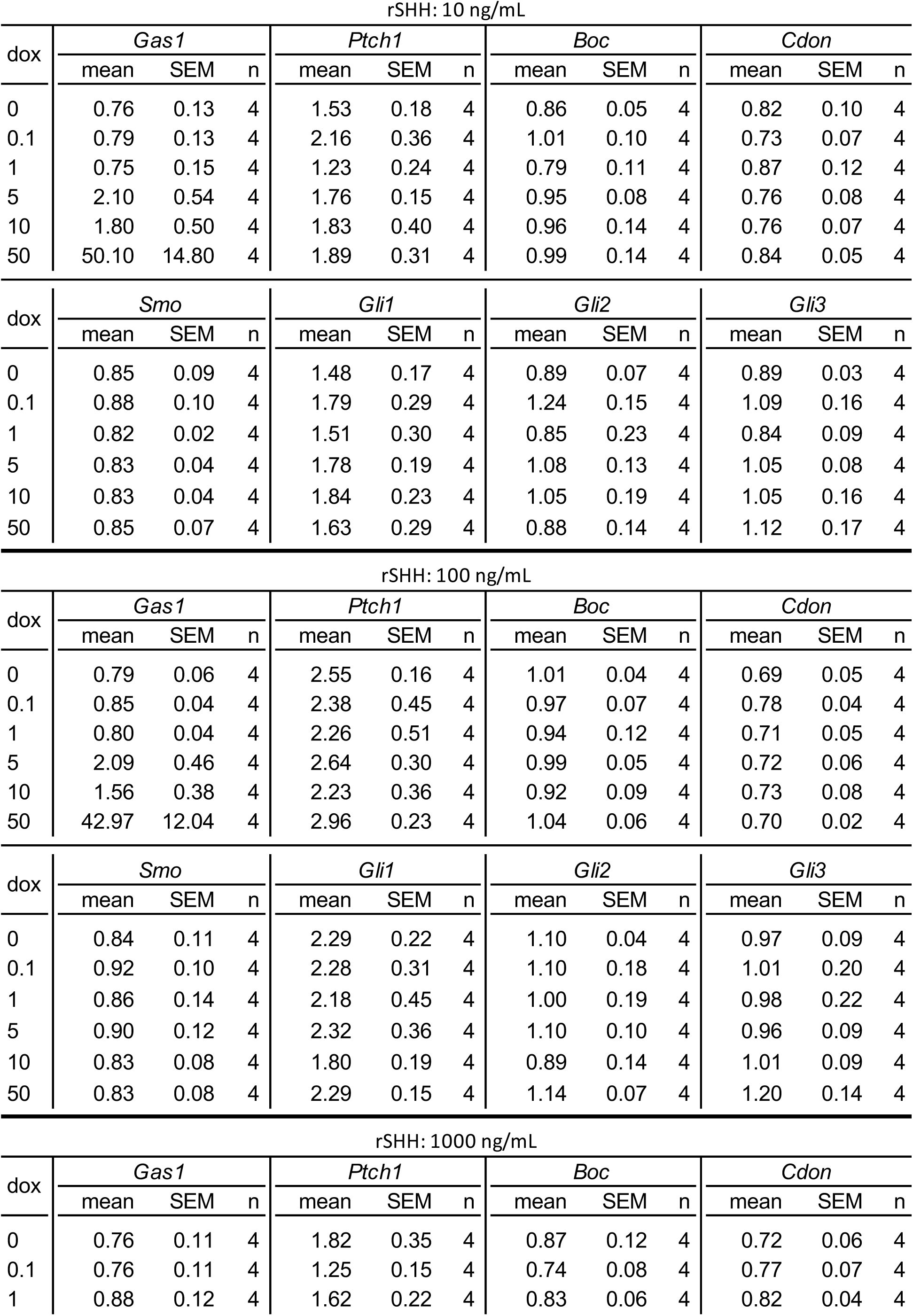
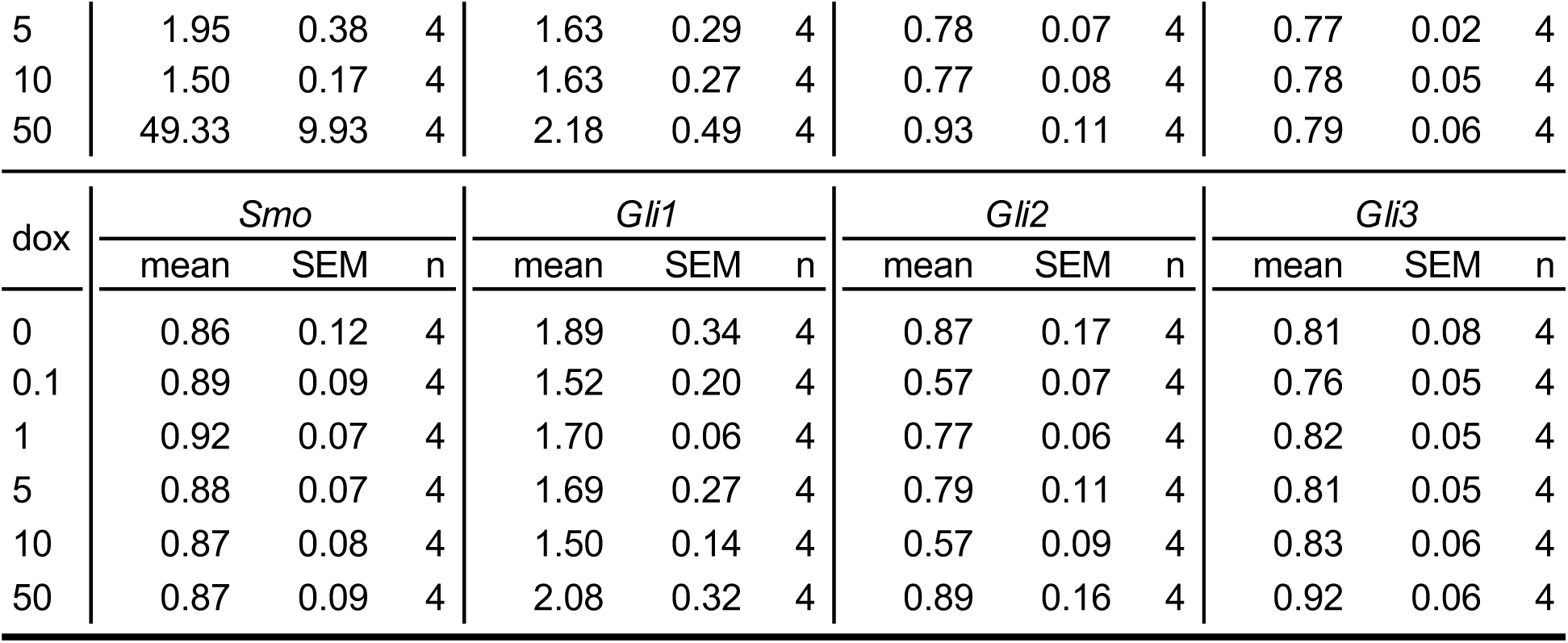
SHH pathway members mRNA levels in DF1 cells with stably integrated dox-inducible *Gas1* construct after 48 hours of doxycycline hyclate (dox) and recombinant SHH (rSHH) treatment.

| rSHH: 0 ng/mL |  |  |  |  |  |  |  |  |  |  |  |  |
| --- | --- | --- | --- | --- | --- | --- | --- | --- | --- | --- | --- | --- |
| dox | *Gas1* |  |  | *Ptch1* |  |  | *Boc* |  |  | *Cdon* |  |  |
|  | mean | SEM | n | mean | SEM | n | mean | SEM | n | mean | SEM | n |
| 0 | 1.00 | 0.04 | 2 | 1.01 | 0.11 | 2 | 1.00 | 0.05 | 2 | 1.00 | 0.07 | 2 |
| 0.1 | 0.97 | 0.08 | 2 | 1.02 | 0.03 | 2 | 1.00 | 0.03 | 2 | 0.91 | 0.10 | 2 |
| 1 | 1.10 | 0.11 | 2 | 0.94 | 0.26 | 2 | 0.95 | 0.15 | 2 | 0.90 | 0.08 | 2 |
| 5 | 2.61 | 0.55 | 2 | 1.16 | 0.12 | 2 | 1.08 | 0.13 | 2 | 0.82 | 0.08 | 2 |
| 10 | 1.64 | 0.57 | 2 | 0.99 | 0.11 | 2 | 0.83 | 0.10 | 2 | 0.64 | 0.17 | 2 |
| 50 | 60.20 | 7.61 | 2 | 1.28 | 0.01 | 2 | 1.04 | 0.21 | 2 | 0.74 | 0.10 | 2 |
| dox | *Smo* |  |  | *Gli1* |  |  | *Gli2* |  |  | *Gli3* |  |  |
|  | mean | SEM | n | mean | SEM | n | mean | SEM | n | mean | SEM | n |
| 0 | 1.00 | 0.06 | 2 | 1.00 | 0.06 | 2 | 1.01 | 0.11 | 2 | 1.00 | 0.08 | 2 |
| 0.1 | 1.00 | 0.00 | 2 | 0.97 | 0.14 | 2 | 1.05 | 0.06 | 2 | 0.90 | 0.05 | 2 |
| 1 | 0.99 | 0.01 | 2 | 0.82 | 0.03 | 2 | 1.05 | 0.38 | 2 | 0.91 | 0.14 | 2 |
| 5 | 0.96 | 0.03 | 2 | 1.04 | 0.01 | 2 | 1.14 | 0.13 | 2 | 0.97 | 0.03 | 2 |
| 10 | 0.76 | 0.20 | 2 | 0.95 | 0.16 | 2 | 1.02 | 0.14 | 2 | 0.97 | 0.23 | 2 |
| 50 | 0.81 | 0.09 | 2 | 1.13 | 0.01 | 2 | 1.05 | 0.02 | 2 | 1.03 | 0.03 | 2 |
| rSHH: 1 ng/mL |  |  |  |  |  |  |  |  |  |  |  |  |
| dox | *Gas1* |  |  | *Ptch1* |  |  | *Boc* |  |  | *Cdon* |  |  |
|  | mean | SEM | n | mean | SEM | n | mean | SEM | n | mean | SEM | n |
| 0 | 0.84 | 0.01 | 2 | 1.29 | 0.14 | 2 | 0.76 | 0.05 | 2 | 0.70 | 0.15 | 2 |
| 0.1 | 0.83 | 0.03 | 2 | 1.26 | 0.15 | 2 | 0.75 | 0.10 | 2 | 0.73 | 0.14 | 2 |
| 1 | 0.91 | 0.16 | 2 | 1.12 | 0.14 | 2 | 0.77 | 0.04 | 2 | 0.85 | 0.18 | 2 |
| 5 | 2.17 | 0.04 | 2 | 1.38 | 0.04 | 2 | 0.80 | 0.04 | 2 | 0.67 | 0.09 | 2 |
| 10 | 1.95 | 0.01 | 2 | 1.55 | 0.00 | 2 | 0.90 | 0.04 | 2 | 0.78 | 0.03 | 2 |
| 50 | 75.86 | 11.80 | 2 | 0.72 | 0.16 | 2 | 0.59 | 0.13 | 2 | 0.85 | 0.06 | 2 |
| dox | *Smo* |  |  | *Gli1* |  |  | *Gli2* |  |  | *Gli3* |  |  |
|  | mean | SEM | n | mean | SEM | n | mean | SEM | n | mean | SEM | n |
| 0 | 0.70 | 0.10 | 2 | 1.30 | 0.00 | 2 | 1.00 | 0.01 | 2 | 0.88 | 0.01 | 2 |
| 0.1 | 0.69 | 0.11 | 2 | 1.20 | 0.17 | 2 | 1.00 | 0.15 | 2 | 0.88 | 0.05 | 2 |
| 1 | 0.65 | 0.08 | 2 | 1.07 | 0.02 | 2 | 0.87 | 0.12 | 2 | 0.82 | 0.04 | 2 |
| 5 | 0.66 | 0.08 | 2 | 1.33 | 0.00 | 2 | 1.04 | 0.06 | 2 | 0.84 | 0.07 | 2 |
| 10 | 0.77 | 0.04 | 2 | 1.53 | 0.03 | 2 | 1.19 | 0.04 | 2 | 0.99 | 0.04 | 2 |
| 50 | 0.75 | 0.00 | 2 | 1.05 | 0.06 | 2 | 0.54 | 0.13 | 2 | 0.77 | 0.09 | 2 |

rSHH: 10 ng/mL
| dox | *Gas1* |  |  | *Ptch1* |  |  | *Boc* |  |  | *Cdon* |  |  |
| --- | --- | --- | --- | --- | --- | --- | --- | --- | --- | --- | --- | --- |
|  | mean | SEM | n | mean | SEM | n | mean | SEM | n | mean | SEM | n |
| 0 | 0.76 | 0.13 | 4 | 1.53 | 0.18 | 4 | 0.86 | 0.05 | 4 | 0.82 | 0.10 | 4 |
| 0.1 | 0.79 | 0.13 | 4 | 2.16 | 0.36 | 4 | 1.01 | 0.10 | 4 | 0.73 | 0.07 | 4 |
| 1 | 0.75 | 0.15 | 4 | 1.23 | 0.24 | 4 | 0.79 | 0.11 | 4 | 0.87 | 0.12 | 4 |
| 5 | 2.10 | 0.54 | 4 | 1.76 | 0.15 | 4 | 0.95 | 0.08 | 4 | 0.76 | 0.08 | 4 |
| 10 | 1.80 | 0.50 | 4 | 1.83 | 0.40 | 4 | 0.96 | 0.14 | 4 | 0.76 | 0.07 | 4 |
| 50 | 50.10 | 14.80 | 4 | 1.89 | 0.31 | 4 | 0.99 | 0.14 | 4 | 0.84 | 0.05 | 4 |

| dox | *Smo* |  |  | *Gli1* |  |  | *Gli2* |  |  | *Gli3* |  |  |
| --- | --- | --- | --- | --- | --- | --- | --- | --- | --- | --- | --- | --- |
|  | mean | SEM | n | mean | SEM | n | mean | SEM | n | mean | SEM | n |
| 0 | 0.85 | 0.09 | 4 | 1.48 | 0.17 | 4 | 0.89 | 0.07 | 4 | 0.89 | 0.03 | 4 |
| 0.1 | 0.88 | 0.10 | 4 | 1.79 | 0.29 | 4 | 1.24 | 0.15 | 4 | 1.09 | 0.16 | 4 |
| 1 | 0.82 | 0.02 | 4 | 1.51 | 0.30 | 4 | 0.85 | 0.23 | 4 | 0.84 | 0.09 | 4 |
| 5 | 0.83 | 0.04 | 4 | 1.78 | 0.19 | 4 | 1.08 | 0.13 | 4 | 1.05 | 0.08 | 4 |
| 10 | 0.83 | 0.04 | 4 | 1.84 | 0.23 | 4 | 1.05 | 0.19 | 4 | 1.05 | 0.16 | 4 |
| 50 | 0.85 | 0.07 | 4 | 1.63 | 0.29 | 4 | 0.88 | 0.14 | 4 | 1.12 | 0.17 | 4 |

rSHH: 100 ng/mL
| dox | *Gas1* |  |  | *Ptch1* |  |  | *Boc* |  |  | *Cdon* |  |  |
| --- | --- | --- | --- | --- | --- | --- | --- | --- | --- | --- | --- | --- |
|  | mean | SEM | n | mean | SEM | n | mean | SEM | n | mean | SEM | n |
| 0 | 0.79 | 0.06 | 4 | 2.55 | 0.16 | 4 | 1.01 | 0.04 | 4 | 0.69 | 0.05 | 4 |
| 0.1 | 0.85 | 0.04 | 4 | 2.38 | 0.45 | 4 | 0.97 | 0.07 | 4 | 0.78 | 0.04 | 4 |
| 1 | 0.80 | 0.04 | 4 | 2.26 | 0.51 | 4 | 0.94 | 0.12 | 4 | 0.71 | 0.05 | 4 |
| 5 | 2.09 | 0.46 | 4 | 2.64 | 0.30 | 4 | 0.99 | 0.05 | 4 | 0.72 | 0.06 | 4 |
| 10 | 1.56 | 0.38 | 4 | 2.23 | 0.36 | 4 | 0.92 | 0.09 | 4 | 0.73 | 0.08 | 4 |
| 50 | 42.97 | 12.04 | 4 | 2.96 | 0.23 | 4 | 1.04 | 0.06 | 4 | 0.70 | 0.02 | 4 |

| dox | *Smo* |  |  | *Gli1* |  |  | *Gli2* |  |  | *Gli3* |  |  |
| --- | --- | --- | --- | --- | --- | --- | --- | --- | --- | --- | --- | --- |
|  | mean | SEM | n | mean | SEM | n | mean | SEM | n | mean | SEM | n |
| 0 | 0.84 | 0.11 | 4 | 2.29 | 0.22 | 4 | 1.10 | 0.04 | 4 | 0.97 | 0.09 | 4 |
| 0.1 | 0.92 | 0.10 | 4 | 2.28 | 0.31 | 4 | 1.10 | 0.18 | 4 | 1.01 | 0.20 | 4 |
| 1 | 0.86 | 0.14 | 4 | 2.18 | 0.45 | 4 | 1.00 | 0.19 | 4 | 0.98 | 0.22 | 4 |
| 5 | 0.90 | 0.12 | 4 | 2.32 | 0.36 | 4 | 1.10 | 0.10 | 4 | 0.96 | 0.09 | 4 |
| 10 | 0.83 | 0.08 | 4 | 1.80 | 0.19 | 4 | 0.89 | 0.14 | 4 | 1.01 | 0.09 | 4 |
| 50 | 0.83 | 0.08 | 4 | 2.29 | 0.15 | 4 | 1.14 | 0.07 | 4 | 1.20 | 0.14 | 4 |

rSHH: 1000 ng/mL
| dox | *Gas1* |  |  | *Ptch1* |  |  | *Boc* |  |  | *Cdon* |  |  |
| --- | --- | --- | --- | --- | --- | --- | --- | --- | --- | --- | --- | --- |
|  | mean | SEM | n | mean | SEM | n | mean | SEM | n | mean | SEM | n |
| 0 | 0.76 | 0.11 | 4 | 1.82 | 0.35 | 4 | 0.87 | 0.12 | 4 | 0.72 | 0.06 | 4 |
| 0.1 | 0.76 | 0.11 | 4 | 1.25 | 0.15 | 4 | 0.74 | 0.08 | 4 | 0.77 | 0.07 | 4 |
| 1 | 0.88 | 0.12 | 4 | 1.62 | 0.22 | 4 | 0.83 | 0.06 | 4 | 0.82 | 0.04 | 4 |
| 5 | 1.95 | 0.38 | 4 | 1.63 | 0.29 | 4 | 0.78 | 0.07 | 4 | 0.77 | 0.02 | 4 |
| 10 | 1.50 | 0.17 | 4 | 1.63 | 0.27 | 4 | 0.77 | 0.08 | 4 | 0.78 | 0.05 | 4 |
| 50 | 49.33 | 9.93 | 4 | 2.18 | 0.49 | 4 | 0.93 | 0.11 | 4 | 0.79 | 0.06 | 4 |

| dox | *Smo* |  |  | *Gli1* |  |  | *Gli2* |  |  | *Gli3* |  |  |
| --- | --- | --- | --- | --- | --- | --- | --- | --- | --- | --- | --- | --- |
|  | mean | SEM | n | mean | SEM | n | mean | SEM | n | mean | SEM | n |
| 0 | 0.86 | 0.12 | 4 | 1.89 | 0.34 | 4 | 0.87 | 0.17 | 4 | 0.81 | 0.08 | 4 |
| 0.1 | 0.89 | 0.09 | 4 | 1.52 | 0.20 | 4 | 0.57 | 0.07 | 4 | 0.76 | 0.05 | 4 |
| 1 | 0.92 | 0.07 | 4 | 1.70 | 0.06 | 4 | 0.77 | 0.06 | 4 | 0.82 | 0.05 | 4 |
| 5 | 0.88 | 0.07 | 4 | 1.69 | 0.27 | 4 | 0.79 | 0.11 | 4 | 0.81 | 0.05 | 4 |
| 10 | 0.87 | 0.08 | 4 | 1.50 | 0.14 | 4 | 0.57 | 0.09 | 4 | 0.83 | 0.06 | 4 |
| 50 | 0.87 | 0.09 | 4 | 2.08 | 0.32 | 4 | 0.89 | 0.16 | 4 | 0.92 | 0.06 | 4 |

**Supplemental Table 18.** P-values for DF1 cell cultures with stably integrated dox-inducible *Gas1* construct after 48 hours of doxycycline hyclate (dox) and/or recombinant SHH (rSHH) protein treatment (significant values highlighted in gray)

| 0 ng/mL |  | rSHH: 0 ng/mL + dox treatment 48h |  |  |  |
| --- | --- | --- | --- | --- | --- |
| vs |  | *Gas1* | *Ptch1* | *Gli1* | *Boc* |
| dox [ng/mL] | 0.1 | > 1.00E+00 | > 1.00E+00 | > 1.00E+00 | > 1.00E+00 |
|  | 1 | > 1.00E+00 | 1.00E+00 | 9.95E-01 | 9.99E-01 |
|  | 5 | 1.00E+00 | 9.99E-01 | 1.00E+00 | 9.87E-01 |
|  | 10 | > 1.00E+00 | > 1.00E+00 | 1.00E+00 | 8.08E-01 |
|  | 50 | < 1.00E-04 | 9.88E-01 | 9.99E-01 | 1.00E+00 |
|  |  | *Cdon* | *Smo* | *Gli2* | *Gli3* |
|  | 0.1 | 9.33E-01 | > 1.00E+00 | 1.00E+00 | 9.87E-01 |
|  | 1 | 9.16E-01 | 1.00E+00 | 1.00E+00 | 9.93E-01 |
|  | 5 | 5.17E-01 | 1.00E+00 | 9.81E-01 | 1.00E+00 |
|  | 10 | 3.78E-02 | 4.88E-01 | > 1.00E+00 | 1.00E+00 |
|  | 50 | 2.04E-01 | 6.85E-01 | 1.00E+00 | 1.00E+00 |
| 0 ng/mL |  | rSHH: 100 ng/mL + dox treatment 48h |  |  |  |
| vs |  | *Gas1* | *Ptch1* | *Gli1* | *Boc* |
| dox [ng/mL] | 0.1 | > 1.00E+00 | 9.93E-01 | > 1.00E+00 | 9.98E-01 |
|  | 1 | > 1.00E+00 | 9.31E-01 | 9.98E-01 | 9.76E-01 |
|  | 5 | 1.00E+00 | 1.00E+00 | 1.00E+00 | 1.00E+00 |
|  | 10 | 1.00E+00 | 9.06E-01 | 4.64E-01 | 9.44E-01 |
|  | 50 | < 1.00E-04 | 7.79E-01 | > 1.00E+00 | 9.99E-01 |
|  |  | *Cdon* | *Smo* | *Gli2* | *Gli3* |
|  | 0.1 | 8.25E-01 | 9.51E-01 | > 1.00E+00 | 9.99E-01 |
|  | 1 | 1.00E+00 | 1.00E+00 | 9.71E-01 | > 1.00E+00 |
|  | 5 | 9.98E-01 | 9.87E-01 | > 1.00E+00 | 1.00E+00 |
|  | 10 | 9.94E-01 | 1.00E+00 | 6.84E-01 | 1.00E+00 |
|  | 50 | 1.00E+00 | 1.00E+00 | 1.00E+00 | 4.47E-01 |

**Supplemental Table 19.** Mandibular mesenchyme population size in duck and quail embryos with *Gas1* over-expression.

| stage | duck *Gas1*-oe |  |  | quail *Gas1*-oe |  |  |
| --- | --- | --- | --- | --- | --- | --- |
|  | mean | SEM | n | mean | SEM | n |
| HH18 | 62888.9 | 23888.9 | 2 | 5222.2 | 1206.98 | 4 |
| HH21 | 131523.81 | 15732.7 | 21 | 29722.25 | 833.35 | 2 |
| HH24 | 408268.52 | 30175.1 | 6 | 106277.78 | 14040 | 4 |

**Supplemental Table 20.** Mandibular primordia incubation times in trypsin-pancreatin solution.

| stage | incubation times [min] |  |  |
| --- | --- | --- | --- |
|  | duck | chick | quail |
| HH18 | 20 | 18 | 15 |
| HH21 | 25 | 20 | 18 |
| HH24 | 30 | 25 | 23 |
| HH27 | 40 | 35 | 30 |

**Supplemental Table 21.**
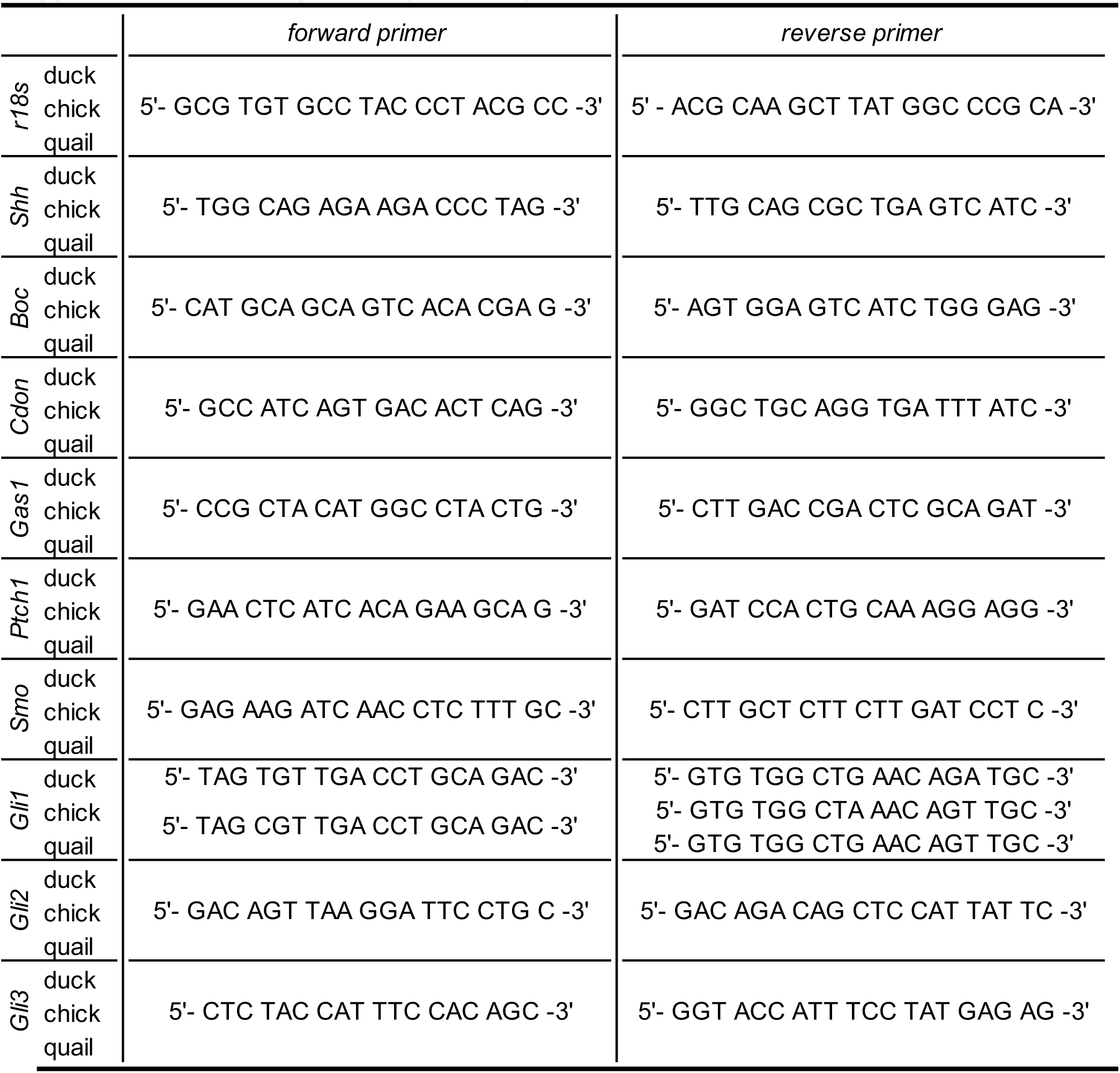
qRT-PCR primer sequences.

**Supplemental Table 22.**
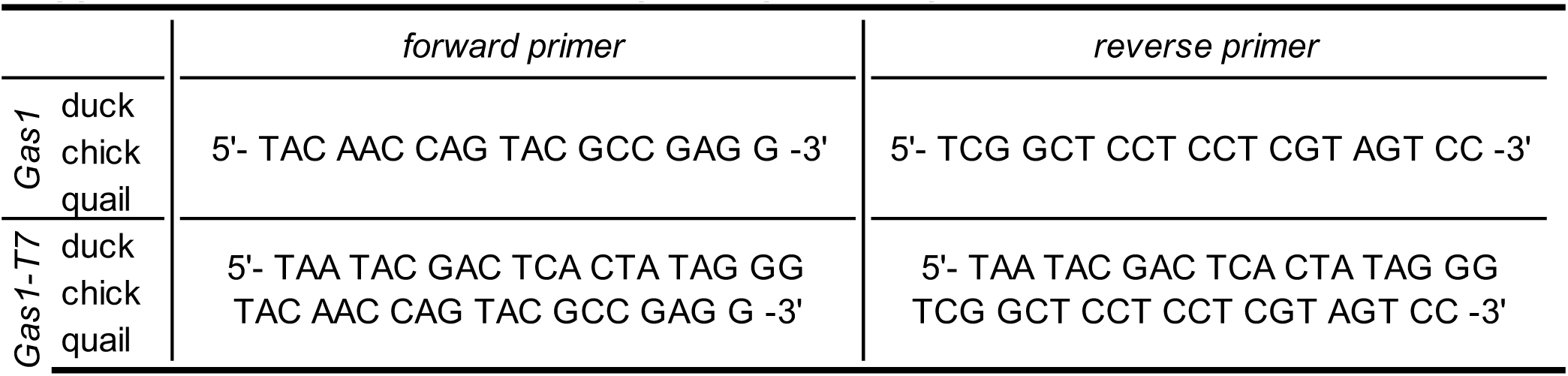
*Gas1* in situs probes primer sequences.

**Supplemental Table 23.**
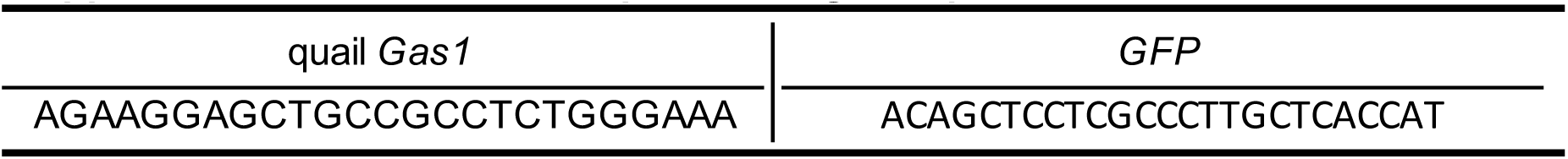
*Vivo-Morpholino oligo* sequences.

